# Acyl chain shortening induced by inhibition of acetyl-CoA carboxylase renders phosphatidylcholine redundant

**DOI:** 10.1101/2021.02.08.429707

**Authors:** Xue Bao, Martijn C. Koorengevel, Marian J.A. Groot Koerkamp, Amir Homavar, Amrah Weijn, Stefan Crielaard, Mike F. Renne, Willie J.C. Geerts, Michal A. Surma, Muriel Mari, Frank C.P. Holstege, Christian Klose, Anton I.P.M. de Kroon

## Abstract

Phosphatidylcholine (PC) is an abundant membrane lipid component in most eukaryotes including yeast. PC has been assigned a multitude of functions in addition to that of building block of the lipid bilayer. Here we show that PC is evolvable essential in yeast by isolating suppressor mutants devoid of PC that exhibit robust growth. The requirement for PC is suppressed by monosomy of chromosome XV, or by a point mutation in the *ACC1* gene encoding acetyl-CoA carboxylase. Although these two genetic adaptations rewire lipid biosynthesis differently, both decrease Acc1 activity thereby reducing the average acyl chain length. Accordingly, soraphen A, a specific inhibitor of Acc1, rescues a yeast mutant with deficient PC synthesis. In the aneuploid suppressor, up-regulation of lipid synthesis is instrumental to accomplish feed-back inhibition of Acc1 by acyl-CoA produced by the fatty acid synthase (FAS). The results show that yeast regulates acyl chain length by fine-tuning the activities of Acc1 and FAS, and indicate that PC evolved by benefitting the maintenance of membrane fluidity.

## INTRODUCTION

The glycerophospholipid phosphatidylcholine (PC) is an essential membrane lipid accounting for at least 50% of total phospholipids in most eukaryotes (van Meer et al., 2008). The exception is presented by several species of green algae that often contain the phosphorus-free betaine lipid diacylglyceryl-*N, N, N*-trimethylhomoserine (DGTS) instead of PC (Giroud and Gerber, 1988; Sato and Furuya, 1985). DGTS and PC both carry a quaternary amine-containing zwitterionic head group and share similar biophysical properties (Sato and Murata, 1991). PC is also present in more than 10% of *Bacteria*, however bacterial PC has not been assigned any essential function (Geiger et al., 2013).

Besides its role as a building block of lipid bilayers, PC has regulatory functions in signal transduction and metabolic regulation in eukaryotes. For example, specific molecular species of PC serve as endogenous ligands for peroxisome proliferator-activated receptor-α (PPARα) and liver receptor homologue 1 (LRH1), respectively (Chakravarthy et al., 2009; Lee et al., 2011). Loss-of-function mutations of key PC biosynthetic enzymes cause a wide spectrum of human pathologies (van der Veen et al., 2017). Furthermore, alterations in PC metabolism have been implicated in cancer (Ridgway, 2013).

By their sheer abundance, PC and its metabolic precursor phosphatidylethanolamine (PE) are important players in determining physical membrane properties such as membrane fluidity and intrinsic curvature that impact the function of membranes and membrane proteins (Covino et al., 2018; de Kroon et al., 2013). Whereas PC has an overall cylindrical molecular shape that makes it ideally suited to build the membrane bilayer matrix, PE is a lipid with non-bilayer propensity that can adopt a conical shape depending on its acyl chain composition (Renne and de Kroon, 2018). The increased PC/PE ratio induced by obesity in mouse liver was found to inhibit Ca^2+^ transport by SERCA, causing ER stress (Fu et al., 2011). A decrease in PC/PE ratio in mouse liver induces steatohepatitis, and ultimately causes liver failure due to loss of membrane integrity (Li et al., 2006).

The tolerance of the model eukaryote *Saccharomyces cerevisiae* towards variation in membrane lipid composition, makes it ideally suited for addressing the functions of lipid classes in membrane lipid homeostasis (de Kroon et al., 2013). The yeast double deletion mutant *cho2opi3* lacking the methyltransferases converting PE to PC, relies on supplementation with choline for the synthesis of PC by the CDP-choline route (Figure 1A) (Kodaki and Yamashita, 1989; Summers et al., 1988), and has been used to manipulate cellular PC content. Both DGTS and phosphatidyldimethylethanolamine (PDME), a lipid containing two instead of three *N*-methyl groups with physical properties similar to PC, can substitute for PC in *cho2opi3* (Boumann et al., 2006; McGraw and Henry, 1989; Riekhof et al., 2014), demonstrating that PC is dispensable for yeast growth.

**Figure 1.**
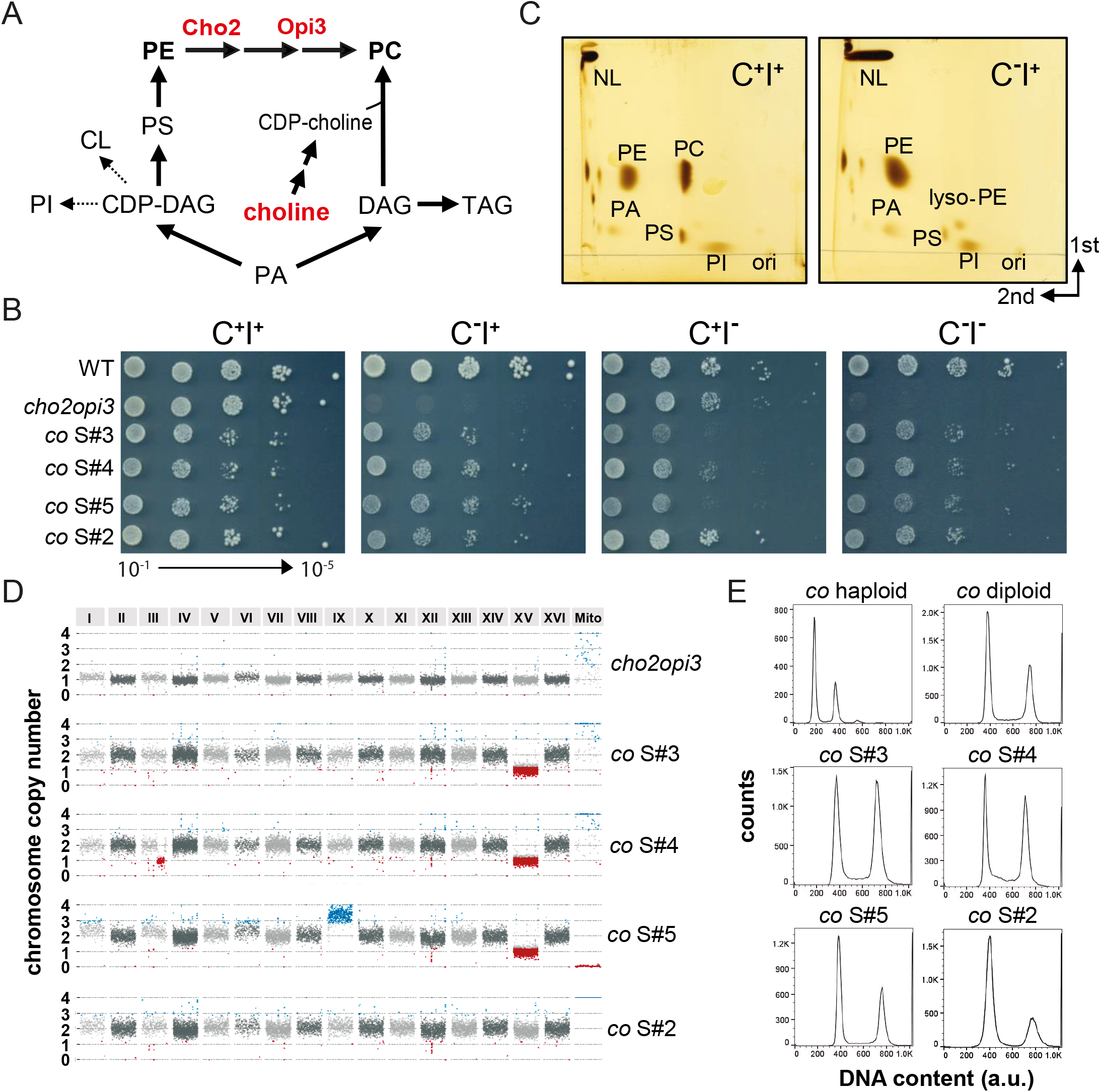
Phenotype and Karyotype of Evolved *cho2opi3* Suppressors. (A) Cartoon depicting the biosynthetic pathways producing PC in yeast. (B) Ten-fold serial dilutions of 1 OD_600_ unit/mL of the indicated strains were spotted on SD plates containing 0 (C^-^) or 1 mM choline (C^+^) and 0 (I^-^) or 75 μM inositol (I^+^) and incubated at 30°C for 3 d. (C) 2D-TLC analysis of total lipid extracts of *co*S#2 cells cultured in SD C^+^I^+^ and C^-^I^+^; ori, origin; NL, neutral lipids. (D) Read-depth analysis indicating monosomy of chromosome XV in *co*S#3, S#4, and S#5, but not in S#2. Each data point represents the median chromosome copy number per 5-kb bin plotted over the genome, with alternating colors for each successive chromosome and the mitochondrial DNA. (E) Representative DNA content profiles of haploid and diploid *cho2opi3* controls (cultured in C^+^) and the indicated *cho2opi3* suppressor strains. See also Figure S1, S2 and Data S1.

Here, we report the isolation and characterization of *cho2opi3* suppressor mutants that exhibit sustained growth in the absence of choline. As the suppressors do not contain PC or a PC substitute, elucidation of the mechanism of suppression provides an unbiased route to address PC function. The choline auxotrophy of *cho2opi3* is suppressed by 2n-1 monosomy of chromosome XV or by a point mutation in the *ACC1* gene encoding acetyl-CoA carboxylase. The genetic changes in both suppressors shorten average acyl chain length due to reduced activity of Acc1. Inhibition of Acc1 is sufficient for suppressing choline auxotrophy as evidenced by the rescue of *cho2opi3* by soraphen A, a specific inhibitor of Acc1. The results indicate that the suppression by chromosome XV monosomy relies on inhibition of Acc1 by accumulating acyl-CoA, providing novel clues about the regulation of acyl chain length by the interplay between Acc1 and the fatty acid synthase complex (FAS). Based on the compensatory changes in the PC-free lipidomes, we propose that the acquisition of PC during evolution provided selective advantage in maintaining membrane physical properties, membrane fluidity in particular.

## RESULTS

### Phenotype and genotype of evolved PC-free yeast *cho2opi3* suppressors

After incubating the *cho2opi3* mutant on choline-free agar plates at 30°C for 14 days, *cho2opi3* suppressor clones were obtained. Most of the clones exhibit sustained growth in the absence of choline, and can be stored as and revived from -80°C glycerol stocks in choline-free SD medium (SD C^-^). A subset of four *cho2opi3* (*co*) suppressor clones, *co*S#2-S#5, was characterized in detail (Fig 1). In contrast to their choline auxotroph *cho2opi3* parent, *co*S#2-#5 grow robustly in the absence of choline, albeit slower than the corresponding WT, irrespective of supplementation with inositol (Fig 1B). The effect of inositol was examined because of its key role in the phosphatidic acid (PA)-mediated transcriptional regulation of phospholipid biosynthesis genes containing UAS_INO_ (Henry et al., 2012). Remarkably, in the absence of inositol, *co*S#3-#5 grow better without than with choline present (Fig 1B), suggesting a choline-sensitive requirement for inositol. The doubling times observed in the corresponding liquid media (Fig S1A) largely recapitulate the growth phenotypes on agar plates.

Analysis by thin layer chromatography (TLC) of total lipid extracts of the suppressors cultured in SD C-indicated that the suppressors are devoid of PC, leaving PE as the predominant membrane lipid (Fig 1C). MS analysis corroborated this result. PC could not be detected in negative ion mode as acetate-adduct, nor in positive ion mode as H^+^-adduct. Fragmentation in the positive ion mode did not reveal the phosphocholine head group.

To elucidate the nature of the adaptation, *co*S#2-#5 were subjected to whole genome sequencing (WGS). WGS did not reveal single nucleotide polymorphisms (SNPs), insertions or deletions shared by the four suppressors (Data S1). However, analysis of chromosome copy number by WGS and fluorescence-activated cell sorting (FACS) revealed changes in ploidy (Fig 1D and E). Suppressors *co*S#3, #4 and #5 exhibit 2n-1 aneuploidy, by losing a copy of chromosome XV (chr XV) after genome duplication. In addition, *co*S#4 lost part of the right arm of one copy of chr III, whereas *co*S#5 gained an extra copy of chr IX and lost its mitochondrial DNA. Ploidy changes, including aneuploidy with gain or loss of chromosomes, are common in adaptive evolution of yeast mutants lacking (non-)essential genes (Liu et al., 2015; Storchova, 2014; Szamecz et al., 2014). Partial karyotype analysis by FACS analysis and a quantitative polymerase chain reaction (qPCR)-based assay (Pavelka et al., 2010) addressing chr XV with chr I, IV, VI and IX as controls, was applied to an extended set of suppressor clones. Like *co*S#3-#5, suppressors *co*S#6-#11 exhibit chr XV monosomy, and similar to *co*S#5, *co*S#8 and S#9 gained extra copies of chr IX (Fig S1B). Generation of (2n-1) suppressors from a diploid *co* strain proceeds more readily than from its haploid counterparts (Figure S1C), suggesting that genome duplication is limiting.

The odd one out is *co*S#2 that turned diploid and retained both copies of chr XV (Fig 1D and E). WGS of *co*S#2 revealed a homozygous point mutation in the *ACC1* gene encoding acetyl-CoA carboxylase, catalyzing the rate limiting step of FA synthesis (Tehlivets et al., 2007). Adenosine at position 657039 of both copies of chr XIV is replaced by cytosine, resulting in the substitution of asparagine at position 1446 of Acc1 by histidine (N1446H; Data S1). Suppressors *co*S#2-S#5 retained the *MATα* mating type as shown by their ability to mate with a threonine-auxotrophic *MAT****a*** strain (Fig S1D), indicating that genome duplication was by endoreduplication rather than mating preceded by mating type switch (Harari et al., 2018).

Previous research showed that propanolamine (Prn) could substitute for choline in supporting growth of *cho2opi3* (Choi et al., 2004). This finding was unexpected, since the physical properties of phosphatidylpropanolamine (PPrn) resemble those of PE rather than PC (Storey et al., 2001). In our hands Prn does not support growth of *cho2opi3* cells, however, suppressors generated on choline-free agar plates supplemented with 1 mM Prn also grow without supplements and exhibit chr XV monosomy (Fig S1E). In retrospect, our data suggest that Choi *et al*. (2004) may have studied *cho2opi3* suppressors.

We conclude that PC biosynthesis is essential in yeast. However, the requirement for PC can be overcome by adaptive evolution.

### Ultrastructure of PC-free yeast

Electron microscopy for morphological examination of PC-free cells revealed that PC-free *co*S#3, S#4 and, to a lesser extent, S#2 (Fig 2E, C, and J) show accumulation of lipid droplets (LD) compared to WT and the *cho2opi3* parent cultured with choline (Fig 2A and B). Quantitation of the area occupied by LD in 2D projection images shows a nearly 3-fold increase in *co*S#3 and S#4 compared to WT and parent (Fig S2A). Other salient features of PC-free cells include the “spikes” of ER often surrounding LD (Fig 2D), in agreement with LD being formed at and staying connected to the ER (Jacquier et al., 2011). In PC-free S#3 and S#2, proliferation of the ER is apparent from protrusions in the nuclear envelope, adopting a “brass-knuckles” shape that occasionally pushes into the vacuole (Fig 2F, I, and J). These structures are reminiscent of the nuclear envelope morphology of a temperature sensitive *acc1* mutant at the restrictive temperature (Schneiter et al., 1996). In S#2 unidentified vacuolar structures accumulate at the limiting membrane of the star-shaped vacuole (Fig 2K). Some mitochondria have aberrant morphology with sheet-like cristae membranes, often detached from the inner boundary membrane (Fig 2G). Given the defects in mitochondrial structure, it is not surprising that *cho2opi3* suppressors do not grow on the non-fermentable carbon source glycerol without choline (Fig S2B). Upon supplementing choline, the PC-free cells return to wild type morphology after 3 doublings (Fig 2H and L), with smaller LD (Fig S2A) often found anchored to the vacuole, suggesting that removal of LD involves lipophagy (van Zutphen et al., 2014).

**Figure 2.**
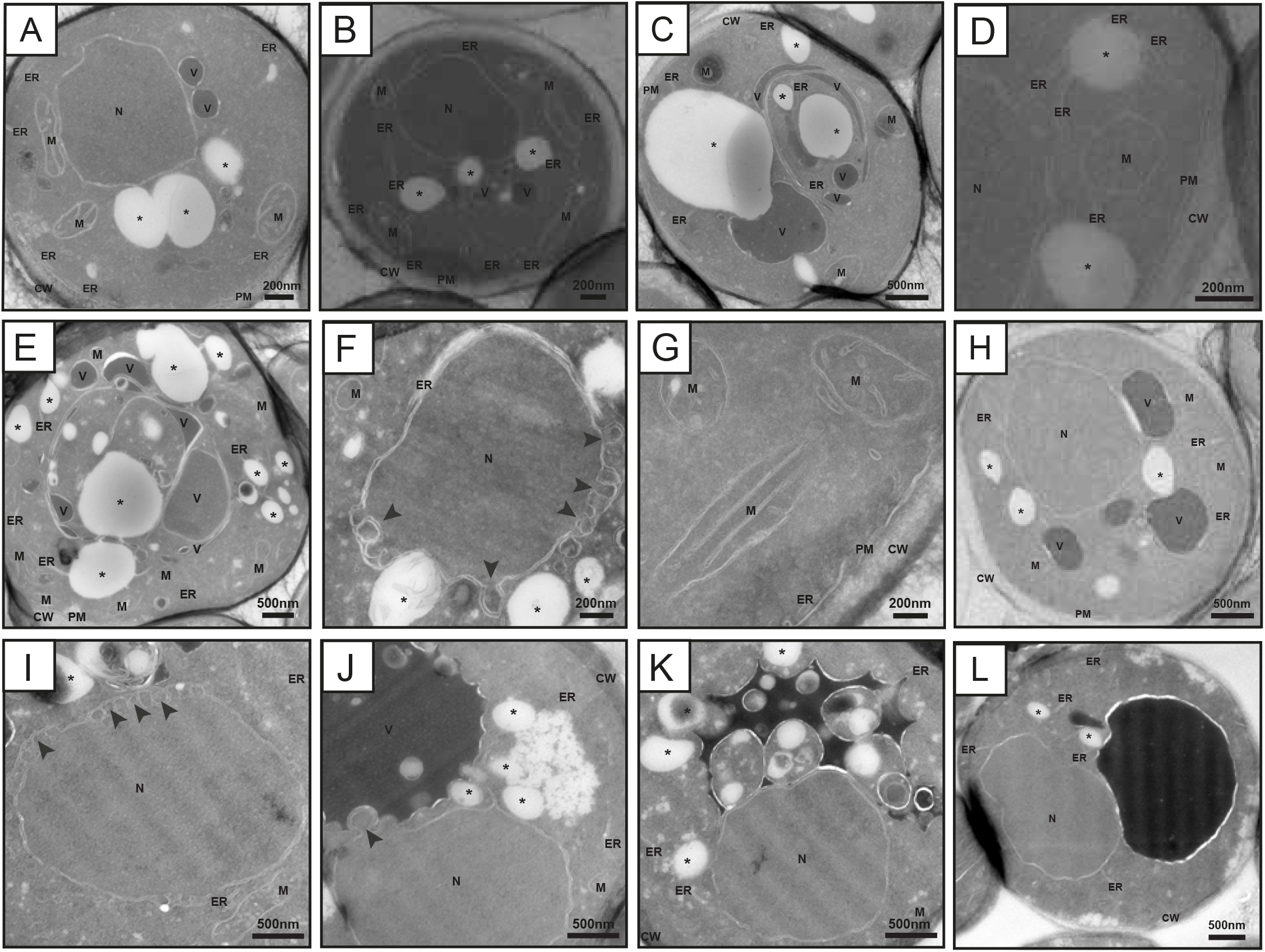
Ultrastructure of PC-free yeast cells. Wild type cultured in C^-^ (A), *cho2opi3* cultured in C^+^ (B), *co*S#4 (C-D), *co*S#3 (E-G), and *co*S#2 (I-K) cultured in C^-^ were analyzed by electron microscopy. In addition, *co*S#3 (H) and *co*S#2 (L) are shown after culture in C^+^ for 3 generations. The arrow heads (F, I, J) point to protrusions in the nuclear envelope. CW, cell wall; PM, plasma membrane; M, mitochondria; N, nucleus; V, vacuole; *, lipid droplet. Scale bars as indicated.

### Monosomy of chromosome XV or a point mutation in *ACC1* is sufficient to suppress choline auxotrophy in a *cho2opi3* mutant

To investigate whether chr XV monosomy is sufficient to suppress choline auxotrophy, a 2n-1 *cho2opi3* strain was constructed by counter selection against a conditionally stable copy of chr XV as described (Reid et al., 2008). Insertion of the *GAL1* promoter and a *URA3* marker adjacent to the centromere (*CEN15*) enabled *CEN* destabilization on galactose-containing medium, and counter selection against *URA3* with 5-fluoroorotic acid (5-FOA), respectively. 5-FOA-induced loss of the destabilized copy of chr XV conferred uracil-auxotrophy while suppressing choline-auxotrophy (Fig 3A), unequivocally demonstrating that chr XV monosomy rescues the choline auxotrophy of *cho2opi3*. In the absence of FOA, suppressors of choline auxotrophy appear more frequently with than without uracil (Fig 2A), as expected based on probability theory. Engineered *co* S(2n-1) and evolved *co*S#3 and S#4 exhibit similar growth phenotypes in the presence or absence of choline and/or inositol (Fig 3B).

**Figure 3.**
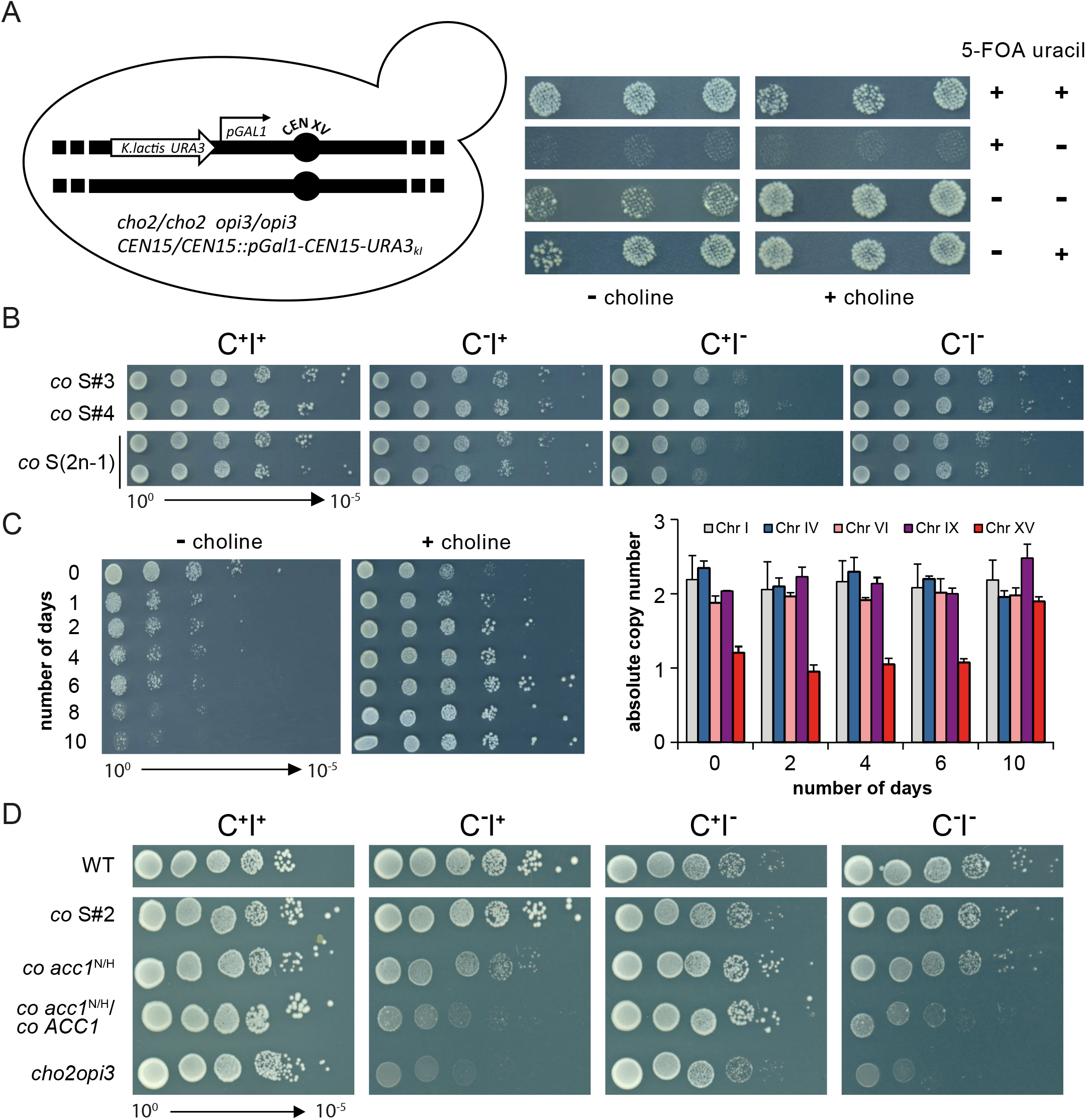
Choline auxotrophy of *cho2opi3* is suppressed by monosomy of chromosome XV or a point mutation in *ACC1*. (A) Induction of chr XV loss in three independent diploid *co/co* clones containing a conditionally stable chr XV suppresses choline auxotrophy. After culture on solid YPGal, cell patches were replica-plated on SD-plates with or without choline, uracil and 5-FOA, as indicated, and incubated at 30 °C for 4 d. (B) Growth of ten-fold serial diluted engineered *co* S(2n-1) and evolved *co* S#3 and S#4 on C^+/-^ I^+/-^ SD at 30 °C for 4 d. (C) Choline supplementation induces endoduplication of chr XV in aneuploid *cho2opi3* suppressors. Growth phenotype on SD C^-^ and C^+^ (4 d at 30 °C) and absolute copy number of chr I, IV, VI, IX and XV as determined by qPCR and FACS after culturing *co* S#4 in SD C^+^ for the number of days indicated with daily passage to fresh medium at OD_600_ 0.05. The error bars represent the variation between assays using primers complementary to non-coding regions on the left and right arm of each chr, respectively. (D) Serial dilutions of the strains indicated were spotted on SD C^+/-^ I^+/-^ and incubated at 30 °C for 4 d.

Upon culture in SD C^+^, 2n-1 suppressors gradually lose the ability to grow in SD C^-^ (and improve growth in SD C^+^) over a period of 6-10 days (Fig 3C). This is accompanied by gain of a second copy of chr XV by endoduplication, in agreement with restoration of euploidy after removal of selection pressure (Chen et al., 2012; Reid et al., 2008).

The N1446H mutation was introduced in Acc1 in the *co* background by CRISPR-Cas9. The engineered haploid *co acc1*^N/H^ mutant recapitulates the growth phenotype of *co*S#2 (Fig 3D), proving that a single point mutation in *ACC1* renders PC redundant. By crossing *co acc1*^N/H^ to *co*, a *co ACC1*/*acc1*^N/H^ heterozygous diploid was generated that shows intermediate growth in SD C^-^.

### PC-free yeast cells accumulate triacylglycerol and exhibit shortening of average acyl chain length

Since Acc1 activity is directly linked to changes in lipid metabolism, we subjected the engineered suppressor strains, parent and WT to mass spectrometry-based shotgun lipidomics (Fig 4, Data S2). After culture in choline-free medium, *co acc1*^N/H^ and *co* S(2n-1) show an almost two-fold increase in membrane lipid content compared to WT and parent strain, accompanied by 3- and 10-fold increases in triacylglycerol (TAG) in *co acc1*^N/H^ and *co* S(2n-1), respectively, reflecting increased FA and glycerolipid synthesis in PC-free suppressors (Fig 4A). Supplementation of choline reduces the level of membrane lipids and TAG in *co acc1*^N/H^ to and below WT level, respectively. In *co* S(2n-1), lipid content is also reduced by choline but stays above WT/parent levels (Fig 4A). Compared to TAG, the ergosterolester content shows a modest increase in the suppressors that in *co* S(2n-1) is not affected by choline (Fig 4A).

**Figure 4.**
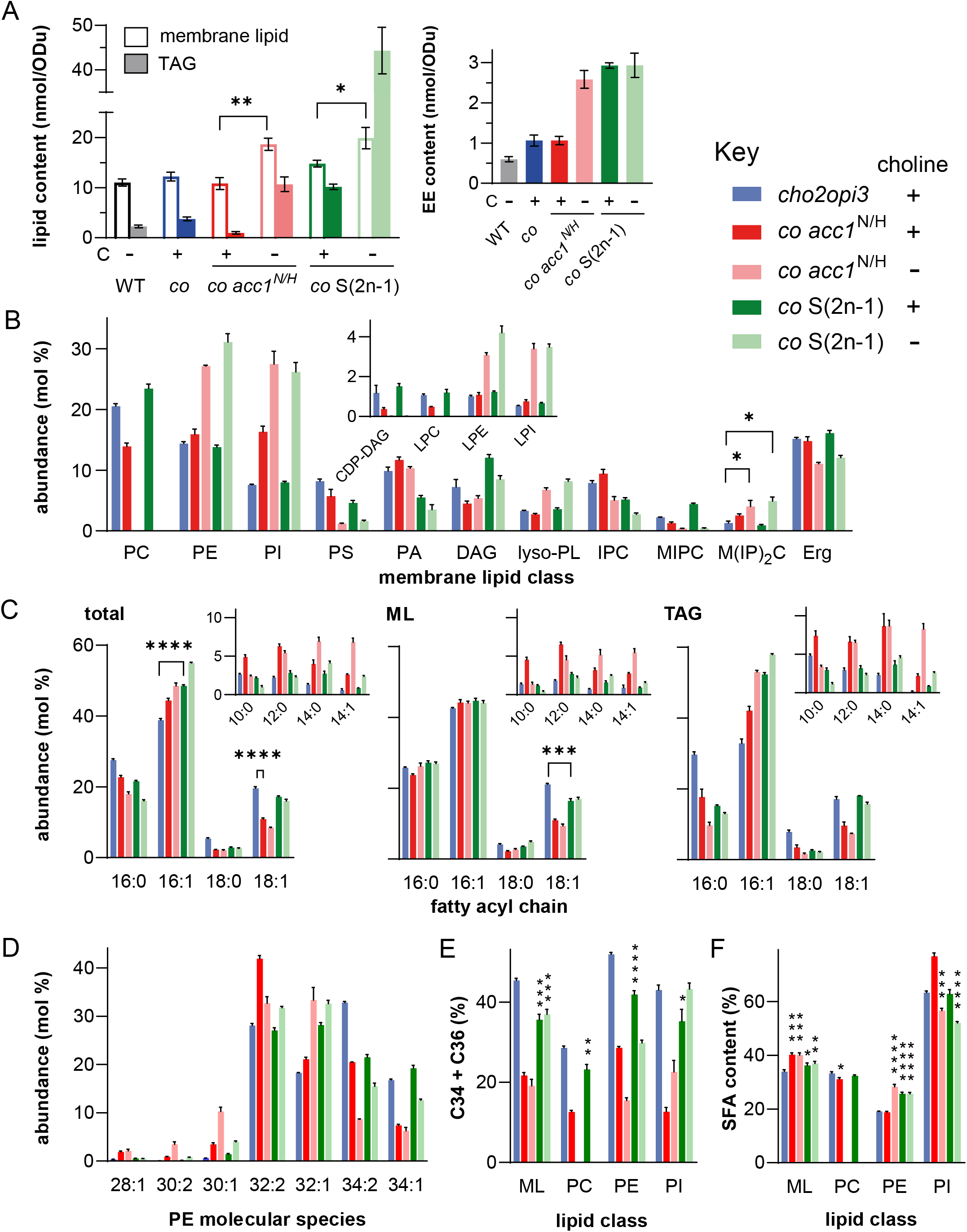
The lipidome of PC-free *cho2opi3* suppressors shows increased lipid content and shortening of average acyl chain length. (A) Membrane lipid and TAG content, and ergosterolester content (EE, inset) per OD_600_ unit of the yeast strains indicated after culture to mid-log phase in SD with or without 1 mM choline (C); * p < 0.05, ** p < 0.01, unpaired two-tailed t-test of the indicated bar compared to the C^+^ condition. (B) Membrane lipid class composition of classes contributing at least 1% of total membrane lipids, the inset shows CDP-DAG and the separate lyso(L)-phospholipids (lyso-PL); (C) Fatty acyl chain profiles of the total lipid, the membrane glycerolipid (ML) and the TAG fraction showing acyl chains that contribute at least 1% of total; (D) PE molecular species profile (sum of carbon atoms in the acyl chains: sum of double bonds in the acyl chains) showing species that contribute at least 2% of total PE; (E-F) Percentage of molecular species containing more than 32 carbon atoms in both acyl chains (C34+C36) (E) and of saturated acyl chains (SFA) (F) in the membrane glycerolipids (ML) and the major membrane lipids; of the indicated strains cultured to mid-log phase in SD C^+/-^. All data were obtained by mass spectrometry and are presented as mean ±SD (n=3); * p < 0.05, ** p < 0.01, *** p < 0.001, **** p < 0.0001, unpaired two-tailed t-test of the indicated bar compared to the *cho2opi3* parent unless indicated otherwise. See also Figure S3, S4 and Data S2.

Lipidomics analysis of the membrane lipid class distribution in *co acc1*^N/H^ and *co* S(2n-1) cultured in SD C^-^ showed that PE and PI take over as major membrane lipids in the absence of PC (Fig 4B). CDP-DAG and PS, the metabolic precursors of membrane lipids and PE, respectively, are depleted, reflecting up-regulated lipid synthesis, in agreement with *cho2opi3* mutants derepressing UAS_INO_ genes in the absence of choline (Boumann et al., 2006; McGraw and Henry, 1989). Derepression of the *INO1* gene, the UAS_INO_ gene with the highest repression/derepression ratio (Henry et al., 2012), results in increased synthesis of inositol that accounts for the increased PI content. The levels of lyso-PE (LPE) and lyso-PI (LPI) rise under choline-free conditions, while the ergosterol (Erg) content is remarkably constant (Fig 4B). Changes in the relative abundance of the sphingolipid M(IP)_2_C relative to its precursors IPC and MIPC follow that of PI (Fig 4B), as reported previously (Jesch et al., 2010). Under choline-replete conditions, membrane lipid composition of *co* S(2n-1) is restored to that of the parent (Fig 4B), that in turn is similar to WT cultured in SD C^-^ (Fig S3A). Suppressor *co* S(2n-1) has lower PI and M(IP)_2_C levels than *co acc1*^N/H^, and like S#3-5 above grows slower in C^+^I^-^ than in C^-^I^-^ (Fig 3B and D), indicating that contrary to *co acc1*^N/H^, 2n-1 suppressors exhibit a choline-sensitive requirement for inositol.

Conventional TLC analysis of phospholipid composition qualitatively confirmed the lipidomics data, and revealed consistent differences between strains when inositol was supplied in the medium except for the PI level remaining largely unchanged (Fig S3B). Possible causes of differences in lipid class levels as determined by MS and TLC, of PI in particular, have been discussed elsewhere (de Kroon, 2017). Phospholipid biosynthesis was examined by pulse labeling *co*S#4 cells with [^32^P]-orthophosphate for 30 min. The reduced percentage of label in PA and the appearance of labeled LPE in the absence of choline (Fig S3C) indicate increased rates of glycerolipid synthesis and PE turnover, respectively.

Analysis of fatty acid content by lipidomics and gas chromatography revealed that both *cho2opi3* suppressors exhibit choline-dependent changes that are hardly affected by supply of inositol (Fig 4C and S3D). Most conspicuously, the relative C18 FA content of *co acc1*^N/H^ is reduced by 50% compared to the parent, an effect observed in both the TAG and the membrane glycerolipid fraction (ML, comprising PC, PE, PI, PS, PA, DAG) with concurrent increases in C10-C14 (Fig 4C). The changes induced by choline deprivation are largely traced back to the TAG fraction with rises in C16:1 at the expense of C16:0 and C18:0 in both suppressors (Fig 4C). The ML of *co acc1*^N/H^ and *co* S(2n-1) shows decreases in C18:1 content of 50 and 20%, respectively, that are compensated for by rises in C10-C14, but otherwise resembles the choline-supplemented parent.

Zooming in on the molecular composition of the individual membrane glycerolipid classes unveils class-and choline-dependent variation between parent and suppressors (Fig 4D and S4A). Both suppressors show increases in PE 32:1 at the expense of PE 34:2 enhanced by choline-deprivation. Moreover, both exhibit a drop in PE 34:1 and rises in C28-32 species, which in *co* S(2n-1) is induced by the absence of choline (Fig 4D). Of note, in choline-free medium a small but significant fraction of PE 34:1 contains C16:1 and C18:0 next to the dominating PE C16:0_C18:1 species (Fig S3E). The shortening of acyl chains, i.e. the decrease in the proportion of C34 and C36 lipids (Fig 4E) that is stronger in *co acc1*^N/H^ than in *co* S(2n-1), is much more pronounced in PE than in PI in PC-free cells. Choline exerts opposite effects on acyl chain length of PE and PI in the suppressors (Fig 4E), by increasing the proportions of C34 in PE and C26-28 in PI (Fig 4D and S4A). The variation in lipid unsaturation is limited, except for increased saturation of PE in *co* S(2n-1) and in PC-free *co acc1*^N/H^ (Fig 4F).

The sphingolipid profiles of *co acc1*^N/H^ are similar to those of the parent strain, whereas *co* S(2n-1) shows an increase in C44 species at the expense of C46, accompanied by an increase in hydroxylation that is stronger and enhanced by choline supply in IPC and MIPC compared to M(IP)_2_C (Fig S4B).

In conclusion, rewiring of lipid synthesis in PC-free suppressors shortens the average acyl chain length and increases the saturation of PE, adaptations consistent with homeostatic control of membrane fluidity and intrinsic curvature (Ernst et al., 2016; Renne and de Kroon, 2018).

### Turnover of PE by the PDAT Lro1 is essential in PC-free *cho2opi3* suppressors

Lro1 is a phospholipid:diacylglycerol acyltransferase that converts DAG to TAG by taking an acyl chain from the *sn*-2 position of glycerophospholipids (Fig 5A) (Dahlqvist et al., 2000; Oelkers et al., 2000), with substrate preference for PE (Dahlqvist et al., 2000; Ghosal et al., 2007; Horvath et al., 2011). The involvement of Lro1 in the rapid turnover of PE into LPE (Fig S3C) and the accumulation of TAG under PC-free conditions (Fig 4A) was investigated. A triple *cho2opi3lro1* deletion mutant does not yield suppressors of choline auxotrophy, in contrast to *cho2opi3* (Fig 5B). Suppressors generated from *cho2opi3lro1* with *LRO1* episomally expressed from the *GAL* promoter lose the ability to grow without choline upon counter selection against the plasmid (Fig 5B and C). Moreover, deletion of *LRO1* in *co acc1*^N/H^ abolishes growth in SD C^-^ (Fig 5D). Galactose-induced expression of Lro1 in *co acc1*^N/H^ *lro1* transformed with p*LRO1* showed that growth and LPE content increase with the galactose concentration in the medium (Fig 5E). In conclusion, Lro1 accounts for the turnover of PE, and is essential in PC-free yeast.

**Figure 5.**
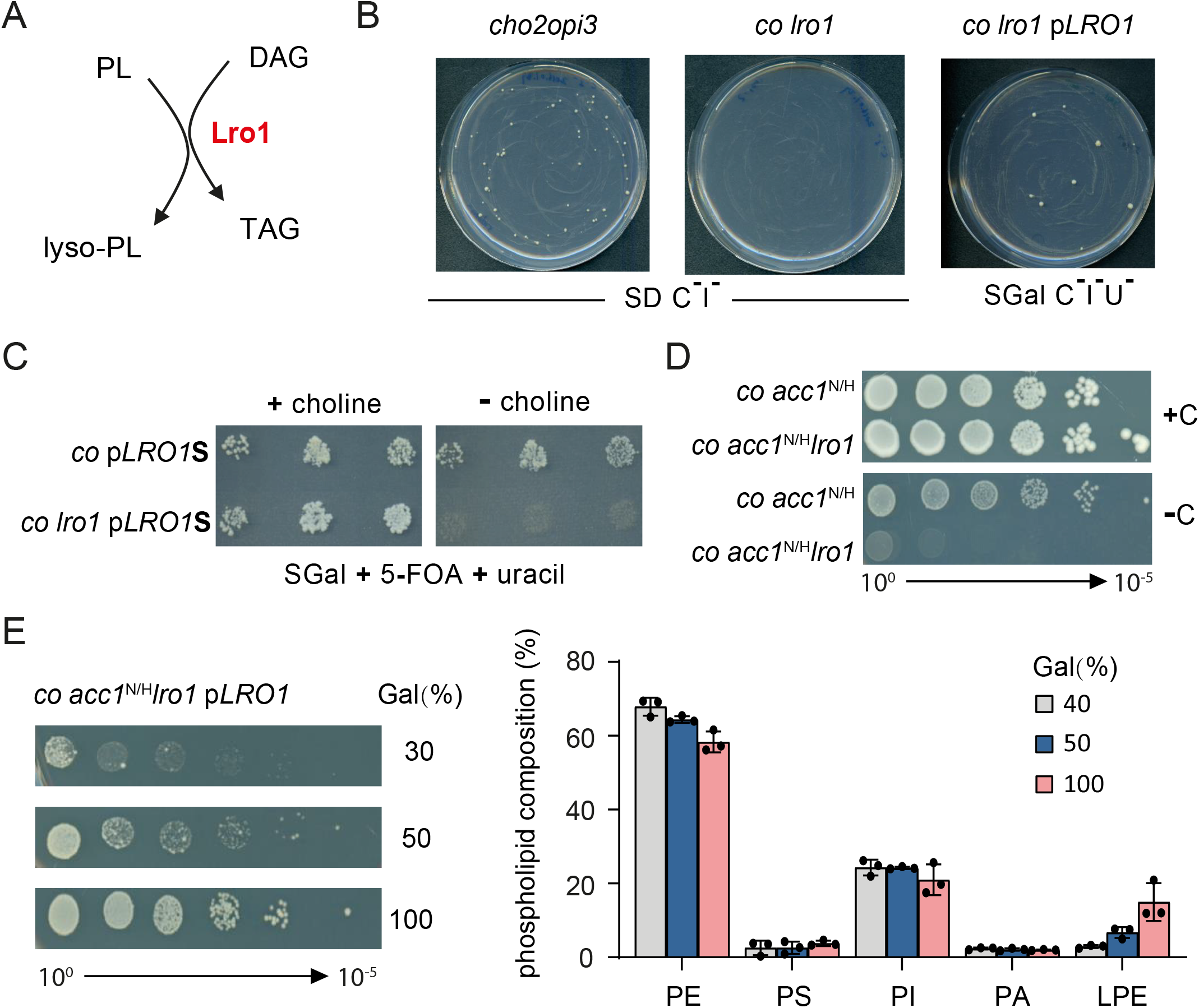
The PDAT Lro1 is essential in *cho2opi3* suppressors. (A) Scheme showing the reaction catalyzed by Lro1. (B) Generation of suppressors of choline auxotrophy of *cho2opi3, cho2opi3lro1*, and *cho2opi3lro1* p*LRO1* on choline-free medium as indicated at 30 °C for 7 d. (C) Three suppressor clones derived of *cho2opi3* and *cho2opi3lro1* transformed with plasmid p*LRO1* were spotted on SGal ura^-^ plates without choline, incubated at 30 °C for 4 d, replica-plated onto SGal C^+^ and C^-^ plates containing 5-FOA and uracil, and incubated at 30 °C for 7 d. (D) Serial dilutions of the strains indicated were spotted on SD C^+/-^ and incubated at 30 °C for 6 d. (E) Growth (30 °C for 6 d) and phospholipid composition of *co acc1*^N/H^ *lro1* p*LRO1* cultured in SD C^-^ ura^-^, containing glucose/galactose mixtures (2%, w/v) as carbon source with the percentage of galactose indicated. Phospholipid composition (±SD, n=3) was analyzed by TLC.

### Inhibition of Acc1 activity by soraphen A abrogates growth of *cho2opi3* suppressors and suppresses choline auxotrophy of the *cho2opi3* parent

Acc1 activity is known to regulate acyl chain length (Hofbauer et al., 2014). The effect on Acc1 activity of the N1446H substitution rescuing choline auxotrophy of *cho2opi3*, was examined by testing sensitivity to soraphen A (SorA), a high affinity (K_d_ 1 nM) inhibitor of Acc1 (Vahlensieck et al., 1994; Weatherly et al., 2004). Already at 0.05 μg/mL, SorA abolishes the growth of *co acc1*^N/H^ and *co*S#2 irrespective of the presence of choline, demonstrating that the point mutation reduces Acc1 activity (Fig 6A).

**Figure 6.**
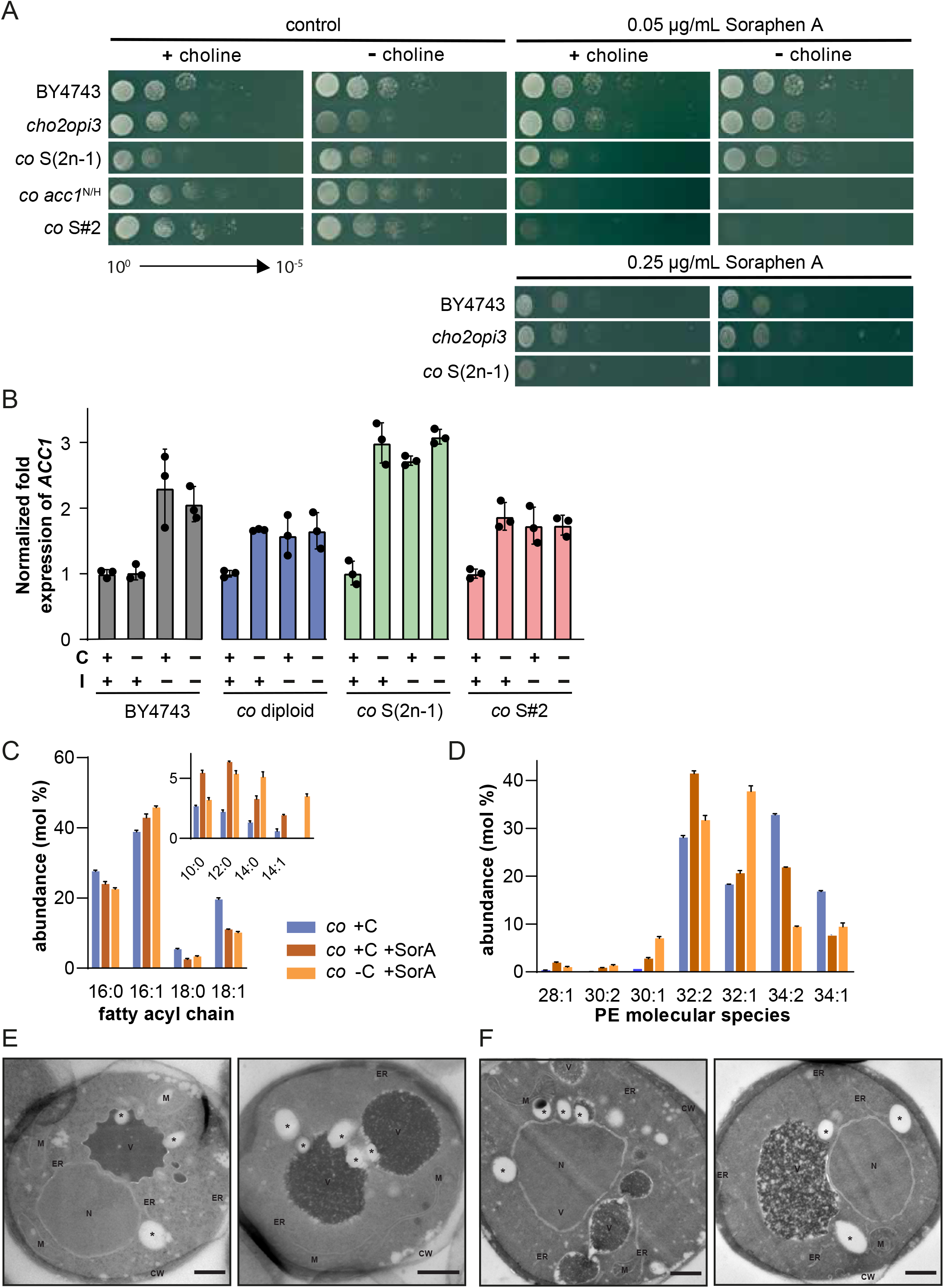
Inhibition of Acc1 activity by soraphen A suppresses choline auxotrophy of the *cho2opi3* parent by reducing average acyl chain length, and abrogates growth of *cho2opi3* suppressors. (A) Ten-fold serial dilutions of 1 OD_600_ unit/mL of the strains indicated were spotted on SD C^+/-^ containing 0.05 or 0.25 μg/mL SorA, and incubated at 30 °C for 3 d. Control plates contain 0.02% (v/v) ethanol. (B) *ACC1* transcript levels after switching the strains indicated from SD C^+^I^+^ to the medium indicated at OD_600_ 0.02 or 0.2 (*co* diploid in C^-^), and culture for 24 h at 30 °C. Data were normalized to *ACT1* and expressed relative to the corresponding strain cultured in C^+^I^+^. The error bars represent SD (n=3). (C-D) Fatty acyl chain profile of the total lipid fraction (C) and PE molecular species profile showing species that contribute at least 1% of total PE (D) of *cho2opi3* transferred to SD C^+^ and SD C^-^ at OD_600_ 0.02 and with 0.05 μg/mL SorA as indicated, and cultured to mid-log phase. Data obtained by mass spectrometry are presented as mean ±SD (n=3). (E-F) EM analysis of *cho2opi3* cells cultured in SD C^+^ (E) and SD C^-^ (F) in the presence of 0.05 μg/mL SorA. The scale bars correspond to 500 nm. See also Figure S5 and Data S2.

Importantly, SorA at 0.05 and 0.25 μg/mL rescues the choline auxotrophy of *cho2opi3*, unequivocally demonstrating that reduced Acc1 activity is sufficient for suppression. In contrast to WT and the parent strain, the *co* S(2n-1) suppressor is sensitive to SorA at 0.25 μg/mL (Fig 6A), which, together with the observed shortening of average acyl chain length (Fig 4C), indicates that reduced Acc1 activity is key to the mechanism of suppression conferred by chr XV monosomy. The mRNA level of *ACC1* goes up 2-3-fold in response to inositol or choline deprivation in both suppressors (Fig 6B) in line with UAS_INO_ regulation of *ACC1* (Chirala, 1992; Henry et al., 2012), and indicating that the lower Acc1 activity in the suppressors is not due to reduced expression.

The lipidome of *cho2opi3* transferred to medium containing 0.05 μg/mL SorA (Data S2) shows a strongly reduced C18 content and a choline deprivation-dependent reduction of C34 species in the PE species profile (Fig 6C and D), similar to *co acc1*^N/H^ (Fig 4C and D). Except for a much lower total lipid content under choline-free conditions (Fig S5A *vs*. 4A), lipidome features of *cho2opi3* in the presence of SorA bear strong resemblance to those of *co acc1*^N/H^ (compare Fig S5B, C, and D with Fig 4B, E, and F). EM analysis shows that the cellular ultrastructure of *cho2opi3* cultured with SorA and choline is indistinguishable from that of choline-deprived cells cultured with SorA (Fig 6E and F), and similar to WT cells (Fig 2A). These data indicate that the morphological changes in the PC-free suppressor strains (Fig 2) are due to a combination of no PC, short acyl chains and lipid overproduction.

Consistent with the essential role of Lro1 under PC-free conditions, SorA does not rescue the choline auxotrophy of *cho2opi3lro1* (Fig S5E). Of note, evolved *co*S#3 and S#4 are more sensitive to SorA than engineered *co* S(2n-1) and *co*S#5 (Fig S5F), indicating that the acquired point mutation in one *ACC1* allele (Data S1) reduces enzyme activity. Accordingly, compared to engineered *co* S(2n-1) the lipidome of *co*S#4 shows less accumulation of TAG and a stronger drop in C18:1 in the absence of choline (Fig S5G and S5H).

Taken together, these results show that the shortening of average acyl chain length induced by inhibiting Acc1 is sufficient for rendering PC redundant.

### mRNA profiling of evolved *cho2opi3* 2n-1 suppressors reveals increased expression of FA-induced ORE genes

As attempts to identify the gene(s) on chr XV requiring lower dosage in *co* S(2n-1) suppressors by complementation with a genomic library failed, whole genome transcript profiling was applied to obtain clues about the mechanism of suppression. The transcriptomes of coS#3, S#4 and S#5 cultured without choline and inositol and that of the *cho2opi3* parent cultured in C^-^I^-^ for up to 3 generations, *i*.*e*. still in log phase (Boumann et al., 2006), were compared to WT (Data S3).

The clustered heatmap (Fig 7A) reveals differences between the transcript profiles of the 3 suppressors that in turn differ dramatically from that of the parent. The transcriptome of the choline-deprived parent shows correlation with the slow growth signature (Fig S6A) that is similar to the environmental stress response (Gasch et al., 2000; O’Duibhir et al., 2014), with increased transcription of stress response genes, accompanied by drops in ribosome biogenesis and amino acid metabolism (Fig 7A, Table S1). In the three suppressors, levels of stress induction are restored to WT (Fig 7A and S6B). Changes in expression of genes governing FA and phospholipid synthesis are modest in parent strain and suppressors (Fig S6B). This also applies to the derepression of UAS_INO_ genes including their most highly regulated representative *INO1*, as expected based on the absence of inositol from the culture medium (Jesch et al., 2006). After preculture in SD C^+^I^+^, deprivation of choline causes a stronger derepression of *INO1* in the *cho2opi3* mutants than deprivation of inositol (Fig S6C), in agreement with previous reports (McGraw and Henry, 1989; Summers et al., 1988). Derepression is stronger in *co*S#2 than in *co* S(2n-1) and *cho2opi3*.

**Figure 7.**
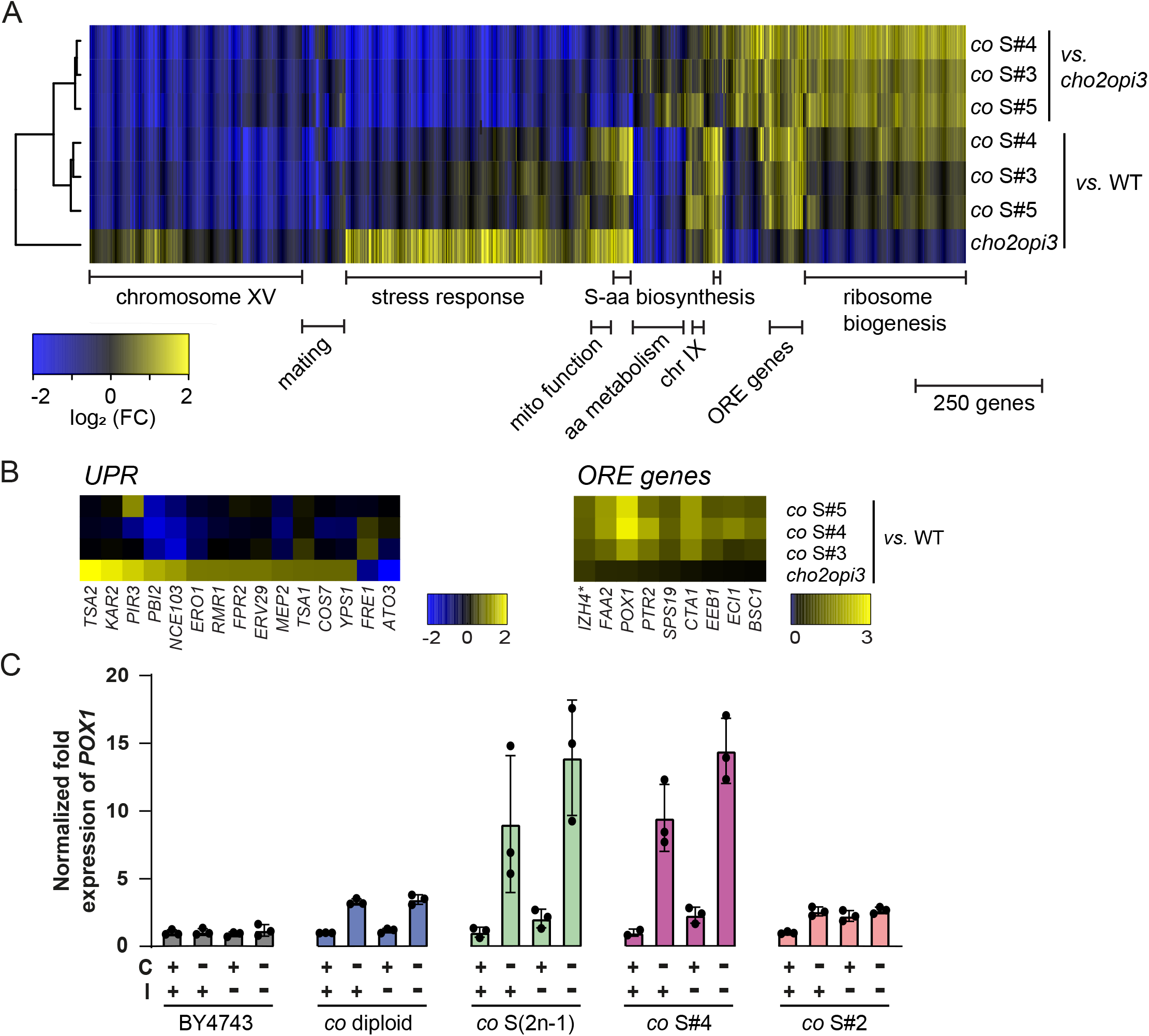
mRNA profiling of evolved *cho2opi3* 2n-1 suppressors reveals upregulation of ORE genes, however β oxidation is not required for suppression of choline auxotrophy. (A) Heatmap showing log_2_ fold changes of mRNA expression in the *cho2opi3* parent strain cultured in SD C^-^I^-^ for 13 h *vs*. wild type, and in *co* S#3, #4, and #5 cultured to mid-log phase in C^-^I^-^ *vs*. wild type and *vs*. parent. Transcripts changing more than 1.7-fold (p < 0.01) in at least one of the comparisons are depicted. Hierarchical clustering was by average linkage (cosine correlation); (functional) categories of enriched transcripts were assigned per cluster. (B) Transcript profiles of the PC-depleted parent and evolved suppressors *vs*. wild type of UPR-induced genes (Kimata et al., 2006; Travers et al., 2000) changing more than 1.7 fold (p < 0.01) in the parent, and of ORE-genes from the cluster in Fig. 6A. Heatmaps show log_2_ fold changes according to the color scales; asterisk (*) indicates gene on chromosome XV. (C) *POX1* transcript levels after switching the strains indicated from SD C^+^I^+^ to the medium indicated at OD_600_ 0.02 or 0.2 (*co* diploid in C^-^), and culture for 24 h at 30 °C. Data were normalized to *ACT1* and expressed relative to the corresponding strain cultured in C^+^I^+^. The error bars represent SD (n=3). See also Figure S6, Table S1 and Data S3.

PC depletion in the parent strain induces the unfolded protein response (UPR, Fig 7B) in agreement with previous reports (Fu et al., 2011; Thibault et al., 2012). Remarkably, the PC-free suppressors do not show strongly increased transcription of UPR-induced genes, including *PBI2* and *PIR3* that are specifically induced by lipid bilayer stress (Ho et al., 2020). Accordingly, RT-qPCR showed modest increases of the well-characterized UPR-induced *KAR2* transcript in WT, *co* S(2n-1) and *co*S#2 upon deprivation of choline, whereas *KAR2* is strongly upregulated in the choline-deprived parent (Fig S6D). Likewise, transcripts related to autophagy upregulated in the choline-deprived parent (Vevea et al., 2015) return to wild type levels in the suppressors (Fig S6B). Since Cho2 and Opi3 are major consumers of *S*-adenosyl methionine (Hickman et al., 2011; Sadhu et al., 2014; Ye et al., 2017), their inactivation impacts the transcription of genes controlling the biosynthesis of sulfur amino acids in both parent and suppressors (Fig 7A and S6B, Table S1).

Zooming in on the transcript levels increased in the evolved 2n-1 suppressors but not in the parent, a cluster enriched in genes containing oleate responsive elements (ORE) was identified, including genes encoding enzymes catalyzing β oxidation (Fig 7A and B, Table S1). ORE genes are induced by the transcription factors Oaf1 and Pip2 upon activation by free FA, most strongly by C16:1 (Karpichev et al., 2008; Phelps et al., 2006). Induction of ORE genes in the presence of the preferred carbon source glucose is unprecedented, raising the question whether β oxidation is required for suppression, e.g. by degrading C18 FA. RT-qPCR confirmed that the transcript of the *POX1* gene encoding acyl-CoA oxidase, the rate limiting enzyme of peroxisomal β oxidation, is upregulated in *co* S(2n-1) and *co*S#4, under choline-free conditions, but not in *co*S#2 (Fig 7C). A *cho2opi3pox1* triple mutant plated on choline-free medium yields suppressors with chr XV monosomy (Fig 7D) like *cho2opi3*, suggesting that the induction of ORE genes in 2n-1 suppressors is a side effect, induced by the high intracellular concentration of free FA.

### Suppression in *co* S(2n-1) depends on inhibition of Acc1 by acyl-CoA

TLC analysis indeed showed that the free FA content of *co* S(2n-1) cultured without choline is increased along with the rise in TAG (Fig S7A). In addition to yielding high levels of free FA and TAG, increased FA synthesis is expected to increase the intracellular concentration of acyl-CoA, a known inhibitor of Acc1 activity (Kamiryo et al., 1976). To test the hypothesis that inhibition of Acc1 activity by acyl-CoA accounts for the suppression of choline auxotrophy in *co* S(2n-1), Dga1 and Gpt2, which consume acyl-CoA in synthesizing TAG and lyso-PA, respectively (Henry et al., 2012), were overexpressed in the suppressors. The ability of *co* S(2n-1) to grow without choline was impaired by overexpressing *DGA1* or *GPT2*, whereas growth of *co acc1*^N/H^ was not affected (Fig 8A), supporting the hypothesis. Moreover, growth of *co* S(2n-1) overexpressing *DGA1* or *GPT2* was restored by SorA, excluding indirect effects on Acc1. Since *DGA1* is localized on chr XV, we verified that loss of 1 copy of *DGA1* is insufficient for suppression of choline auxotrophy (Fig S7B). In conclusion, feed-back inhibition of Acc1 by the product of FAS is crucial for sustained growth of PC-free *co* S(2n-1).

**Figure 8.**
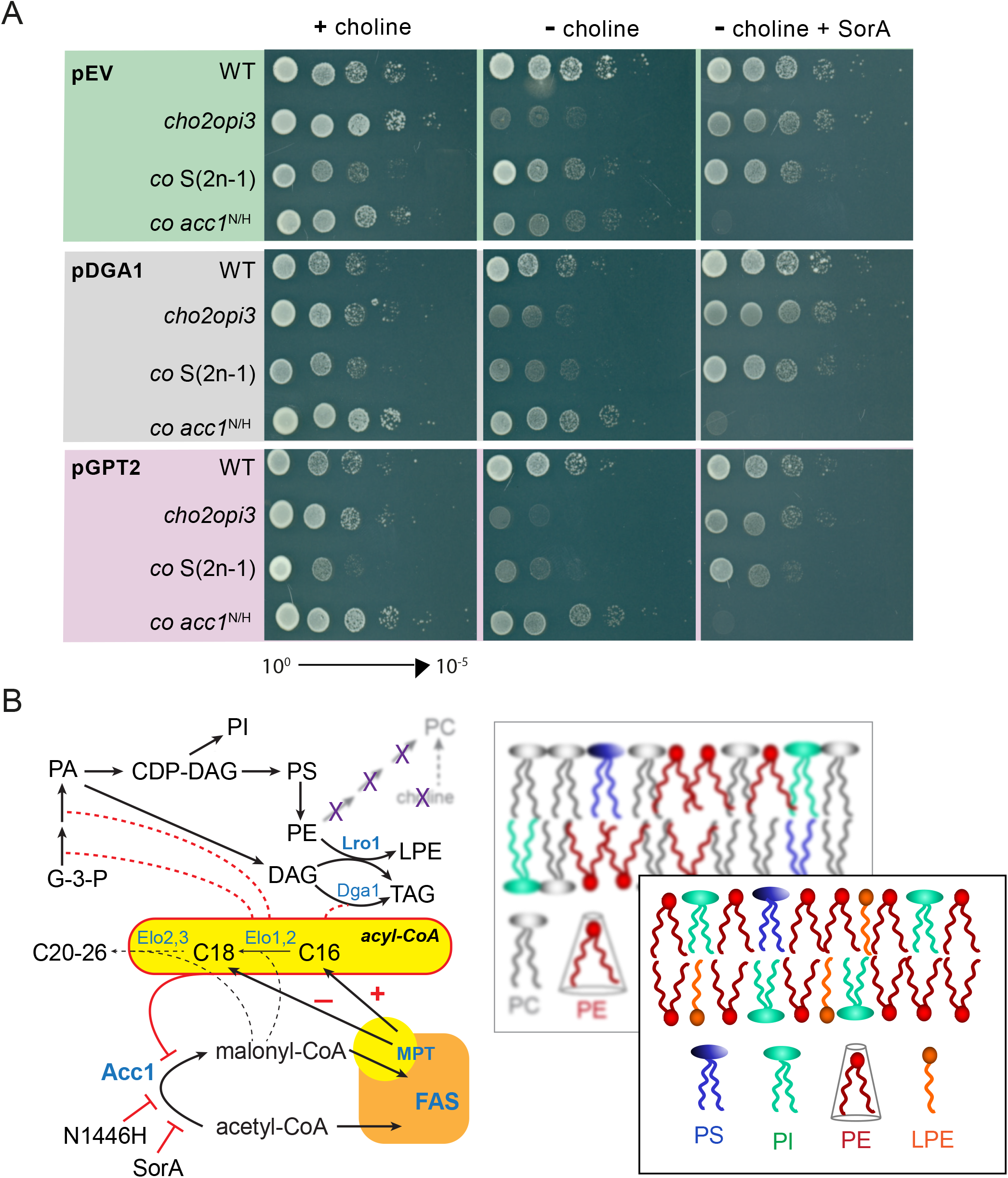
Suppression in 2n-1 suppressors depends on inhibition of Acc1 by an increased concentration of acyl-CoA. (A) Growth of *co* S(2n-1) in the absence of choline is lost when *GPT2* or *DGA1* is overexpressed, and restored by SorA. Plasmids pHEYg-1-*DGA1*, pHEYg-1-*GPT2* and the empty vector control (pEV) were transformed into the strains indicated. After preculture in SD C^+^, ten-fold serial dilutions were spotted on SD C^+/-^ with or without 0.1 μg/mL SorA and incubated at 30 °C for 3 d. (B) Model depicting glycerolipid synthesis and membrane lipid composition in *cho2opi3* suppressor strains when PC has become obsolete (blurred), highlighting (in red) the inhibition of Acc1 that is essential for shortening average acyl chain length that in turn is required for maintaining the physical properties of membranes faced with excess PE. The decreased non-bilayer propensity of PE is indicated by its reduced conical shape. See text for details. See also Figure S7.

## DISCUSSION

### PC biosynthesis is evolvable essential in yeast

The choline auxotrophy of *cho2opi3*, i.e. the requirement for PC, is suppressed by reduced activity of Acc1, as shown by the *acc1*^N/H^ and 2n-1 aneuploid suppressor mutants, and by the rescue by SorA (Fig 8B). The sustained growth of the PC-free suppressors implies that PC or PC substitutes, such as DGTS or PDME, their biosynthesis and turnover do not fulfill essential functions during mitotic fermentative growth of yeast. In contrast, growth on non-fermentable carbon source is abolished, indicating that mitochondrial function is incompatible with the absence of PC (Griac et al., 1996), and/or incompatible with the adaptation. These findings beg new experiments in a PC-free background to address processes in which PC or PC metabolism has been implicated, such as intracellular vesicle trafficking (Bankaitis et al., 2010), sporulation (Mendonsa and Engebrecht, 2009), and mRNA localization (Hermesh et al., 2014).

Our findings qualify PC biosynthesis as an evolvable essential process (Liu et al., 2015). As frequently observed in suppression of severe growth defects due to genetic or environmental perturbations (Chen et al., 2012; Liu et al., 2015; van Leeuwen et al., 2020), a change in ploidy is the prevalent genetic adaptation suppressing PC deficiency. The stress caused by PC depletion probably induces chromosome instability resulting in aneuploidy. Aneuploidy drives rapid adaptation by changing the stoichiometry of gene(s) and by further increasing chromosome instability, imparting fitness gains under adverse conditions (Liu et al., 2015; Pavelka et al., 2010; Torres et al., 2007; Zhu et al., 2012). The phenotypic variation introduced by aneuploidy is immediately clear from the different transcript profiles of *co*S#3-5, while the extra copies of chr IX in some suppressors underscore chromosome instability. After genome doubling, both *co*S#3 and S#4 acquired point mutations in one allele of *ACC1* that reduce Acc1 activity, illustrating that aneuploidy facilitates genetic adaptation (Szamecz et al., 2014; Yona et al., 2012). The *acc1*^N1446H^ mutation in suppressor *co*S#2 preceded genome duplication, suggesting a fitness advantage of the diploid over the haploid PC-free state (*cf*.(Harari et al., 2018)).

### Inhibition of Acc1 is crucial for suppression

Yeast Acc1 is a 500-kDa homodimer, of which the crystal structure has been solved (Wei and Tong, 2015). Asparagine 1446 that is mutated to histidine in *co*S#2, is one of the few conserved amino acids in the Acc1 central (AC) region, and located close to the catalytic carboxyltransfer (CT) domain. The AC region is thought to properly position the biotin carboxylase (BC) and CT dimers for catalysis (Wei and Tong, 2015). We speculate that subtle changes in positioning caused by the N/H substitution account for the reduced activity of Acc1^N1446H^. Acc1 activity is rate limiting fatty acid synthesis, affecting both amount and length of the acyl-CoA’s produced by FAS (consisting of six Fas1 and six Fas2 subunits). Higher concentrations of malonyl-CoA result in the synthesis of longer acyl-CoA’s *in vitro* (Hori et al., 1987; Lynen et al., 1964). Accordingly, the hyperactive Acc1^S1157A^ mutant lacking a phosphorylation site for the Snf1 kinase displays a shift to longer average acyl chain length, along with increased FA synthesis and TAG accumulation (Hofbauer et al., 2014). Conversely, conditional *acc1* mutants exhibit diminished average acyl chain length under restrictive conditions (Schneiter et al., 1996, 2000), similar to *co acc1*^N/H^. Since the sphingolipid species profiles are similar between *co acc1*^N/H^ and parent, the supply of malonyl-CoA probably does not limit the activities of the acyl-CoA elongases Elo1, 2, 3 (Tehlivets et al., 2007).

It is important to realize that *ACC1* and *FAS1/2* are UAS_INO_ genes co-regulated at the transcriptional level by the ER-associated transcription factor Opi1 that senses changes in PA concentration. As the PA level drops, Opi1 translocates into the nucleus to repress the Ino2/4 transcriptional activator complex (Henry et al., 2012). Under choline-free conditions, the derepression of UAS_INO_ genes in *cho2opi3* strains is much stronger than in inositol-deprived WT, which is attributed to the enhanced binding of Opi1 to PA in membranes containing increasing levels of PE (Putta et al., 2016; Young et al., 2010), and to PA with shorter acyl chains (Hofbauer et al., 2014). The derepressed state of UAS_INO_ genes likely explains the increased lipid production in PC-free *co acc1*^N/H^, and contributes to that in *co* S(2n-1). Derepression in *co acc1*^N/H^ is stronger than in *co* S(2n-1), which in addition to shorter average acyl chain length, may be due to haploinsufficiency of *INO4* in the latter. Similarly, *INO4* haploinsufficiency may account for the choline-sensitive inositol requirement of *co* S(2n-1), induced by the CDP-choline pathway depleting PA levels via DAG (Gaspar et al., 2017).

In the PC-free aneuploid *co* S(2n-1) suppressor, the shift to shorter average acyl chain length and the increased sensitivity to SorA are modest, whereas FAS activity is exceedingly high, compared to *co acc1*^N/H^. In fact, the reduced length of sphingolipids in *co* S(2n-1) may be explained by FAS activity competing with the elongase Elo3 for malonyl-CoA (Al-Feel et al., 2003). We propose that the increased FA synthesis conferred by chr XV monosomy is required for inhibition of Acc1 by acyl-CoA (Fig 8B), as depletion of acyl-CoA by overexpression of Dga1 or Gpt2 abolishes suppression. These results identify acyl-CoA as feedback regulator of Acc1 *in vivo*. Inhibition of yeast and mammalian Acc1 by acyl-CoA was previously demonstrated *ex vivo* and *in vitro*, respectively (Kamiryo et al., 1976; Ogiwara et al., 1978). Importantly, the results show that the ratio of Acc1-to-FAS activity rather than Acc1 activity *per se* determines acyl chain length, in agreement with loading of malonyl-CoA competing with release of the mature acyl-CoA at the malonyl/palmitoyl transferase (MPT) domain of FAS, a concept based on *in vitro* studies (Heil et al., 2019). Whether the aneuploidy-induced reduction of the Acc1-to-FAS activity ratio is sufficient to account for suppression in *co* S(2n-1), awaits identification of the haploinsufficient genes on chr XV (likely 2 or more) responsible for the suppression.

### The adaptation obeys the biophysical interplay of phospholipid class and acyl chain composition

The overall shift to shorter acyl chains in the PC-free suppressors compensates for the decrease in membrane fluidity conferred by increased PE content (Dawaliby et al., 2016; Renne and de Kroon, 2018). Concomitantly, it keeps membrane intrinsic curvature in check by reducing PE’s non-bilayer propensity (Fig 8B) (de Kroon et al., 2013). Indeed, responding to increased negative intrinsic curvature by decreasing unsaturation would have aggravated the drop in membrane fluidity, while enhancing acyl chain unsaturation in response to decreasing membrane fluidity would have jeopardized membrane integrity by increasing the non-bilayer propensity of PE. In agreement with this latter notion, rising PE levels do not affect the Mga2 sensor that activates transcription of *OLE1* encoding the yeast desaturase (Ballweg et al., 2020).

The average acyl chain length of PE in the suppressors is reduced by choline-starvation, which is offset by increased acyl chain length of PI. Similar changes in PE profile occur during PC depletion for four generations in the parent strain (Boumann et al., 2006), and presumably proceed by a similar mechanism. The relatively mild UPR induction in PC-free cells indicates that the physicochemical properties of the adapted PC-free membranes are close to wild type (Halbleib et al., 2017).

### PC function and evolution

PC biosynthesis controls the level of PE directly by *N*-methylation and indirectly by shunting PA via DAG into the CDP-choline pathway (Fig 8B). Choline-deprived single *cho2* and *opi3* mutants show a rise in TAG (Fei et al., 2011; Thibault et al., 2012), and growth of a *cho2lro1dga1* mutant in the absence of choline is severely compromised (Garbarino et al., 2009; Vevea et al., 2015), indicating that TAG synthesis is required to buffer defective PC synthesis. Under PC-free conditions, Lro1 becomes essential by degrading PE, compensating for the lack of PC biosynthesis. Whether Lro1 degrades specific PE molecular species, whether its activity is regulated, and whether the LPE produced is just a metabolic intermediate or contributes to bilayer stability by virtue of its more cylindrical molecular shape (Tilcock et al., 1986), are important questions for future research, as is the metabolic fate of LPE.

Bacteria with a high content of unsaturated acyl chains often contain PC (Goldfine, 1984). Inactivation of PC biosynthesis abolishes their growth at higher temperatures, arguing that PC evolved to neutralize the tendency of unsaturated PE to adopt non-bilayer structure (Geiger et al., 2013). In the reverse evolution of *cho2opi3* yeast reported here, the lack of PC is compensated for by acyl chain shortening, in line with PC restraining negative intrinsic curvature conferred by PE. The nature of the compensatory response, *i*.*e*. shortening rather than decreased unsaturation of acyl chains, argues that PC became crucial in maintaining fluidity of eukaryotic membranes during evolution. The temperature-sensitive growth of the PC-free suppressors (Fig S7C) supports this notion.

The PC-free *cho2opi3* suppressor strains add a new model system for use in research aimed at understanding membrane lipid homeostasis and lipid function.

## MATERIALS AND METHODS

### Strains and media

All yeast strains used in this study are listed in the Reagents and Tools Table. Synthetic defined medium (Griac et al., 1996) was supplemented with or without 1 mM choline (C) as indicated. Inositol (I) was supplemented at 75 μM inositol where indicated. 2% (w/v) glucose (SD), 3% (v/v) glycerol (SG), 2% (w/v) galactose (SGal) or mixtures of glucose and galactose were added as carbon source as indicated. YPD and YPGal medium contained 10 g/L yeast extract, 20 g/L bacto-peptone and 20 g/L glucose and galactose, respectively. Solid media contained 2% (w/v) choline- and inositol-free agar (Sigma-Aldrich). 3-amino-1-propanol (Prn) was added from a 1 M stock solution in water adjusted to pH 7.4, to a final concentration of 1 mM. Soraphen A was dissolved in ethanol (2 mg/mL) and added at the concentrations indicated.

### Yeast culture conditions

Strains were cultured at 30 °C unless indicated otherwise. Optical density at 600 nm (OD_600_) was measured with a Hitachi U-2000 double-beam spectrophotometer. The WT strain was pre-cultured in SD without choline (C^-^), whereas *cho2opi3* and derived triple mutants were pre-cultured in SD with choline (C^+^) to the mid-log phase of growth (OD_600_ ∼0.45-1.2). Cells transformed with plasmids were grown in selective SD drop-out media. To deplete phosphatidylcholine, *cho2opi3* cells were collected by filtration or centrifugation as indicated, washed thoroughly with pre-warmed SD C^-^ (30 °C), and transferred to fresh SD C^-^ at the initial OD_600_ indicated (Boumann et al., 2006). SH80 and SH85 were precultured in YPD. Doubling time (DT) was determined based on OD_600_ values at different time points according to:

DT = ln2/μ_max_ with μ_max_ the growth rate during exponential growth, μ_max_ = (lnOD_600_^t2^ -lnOD_600_^t1^)/(t_2_-t_1_).

### Isolation and culture of *cho2opi3* suppressors

The *cho2opi3* double mutant was pre-cultured in SD C^+^ to late log phase (OD_600_ ∼1.5), washed twice with sterile water and resuspended in water at OD_600_ 1. 100 μL of the suspension was spread on a choline-free SD plate. After 10 days at 30 °C,10 *cho2opi3* suppressor strains, *cho2opi3* S#2 to S#11, were obtained. After double confirmation of growth on C^-^ plates at 30 °C, the suppressor strains were frozen as glycerol stocks in SD C^-^. *cho2opi3*S strains were maintained in SD C^-^ medium.

### Adaptation of *cho2opi3* suppressors to choline-containing medium

After pre-culture in choline-free SD to OD_600_∼1, *cho2opi3* S#4 was transferred to SD medium with choline (C^+^) at 30 °C with daily transfer to fresh medium for up to 40 days. Samples were taken every day and frozen as glycerol stocks. After collecting all glycerol stocks, cells were streaked on C^+^, and incubated at 30 °C for 3 days. Single colonies were inoculated in C^+^ liquid medium and tested as indicated.

### Growth phenotype

Cells were pre-cultured to mid-log phase (OD_600_ ∼0.45-1.2) in SD as above, harvested by centrifugation and washed twice with sterile MQ water. The cells were adjusted to OD_600_ 1.0, and serially diluted in 10-fold increments to 10^−5^. 8 μL aliquots of each dilution were spotted onto agar plates containing the medium and supplements indicated. Plates were incubated at 30 °C for the number of days indicated.

### Strain construction

Microbiological techniques followed standard procedures. Diploid *cho2opi3* (*co*) strains were obtained by crossing *co MAT****a*** with *co MATα* obtained by deleting the *OPI3* gene in *cho2* strains as described (Boumann et al., 2004), and selection on lys^-^ met^-^ dropout medium. The *LRO1, DGA1* and *POX1* genes were deleted by standard PCR-based homologous recombination replacing the respective ORFs with the *Sp_HIS3* cassette from the plasmid pFA6a-*HIS3MX6* (Sikorski and Hieter, 1989), using the primers listed in Table S2. Correct integration was verified by colony PCR using primers A and D flanking the regions of *LRO1, DGA1* and *POX1* homology, and primers B and C internal to the *Sp_HIS3* coding region (Table S2).

The genomic single-nucleotide mutation *acc1*^*N1446H*^ was introduced by CRISPR/Cas9 according to published methods (Mans et al., 2015). The pMEL16 backbone containing the 2μ origin of replication and *HIS3* marker was amplified by PCR using conditions and primers as described (Mans et al., 2015). The gRNA insert primers listed in Table S2 were designed with the Yeastriction tool (yeastriction.tnw.tudelft.nl). To obtain the double strand gRNA insert, the complementary gRNA insert primers were dissolved in water to a final concentration of 100 μM, mixed in a 1:1 volume ratio, heated at 95 °C for 5 min, and cooled down to room temperature. The complementary 120 bp repair DNA primers (Table S2) were designed to replace adenosine (A) at position 657039 of chromosome XIV with cytosine (C), and converted to double stranded repair DNA as above. BY4742 and *co MATα* transformed with pCRCT, were co-transformed with 100 ng linearized pMEL16 backbone, 300 ng dsgRNA and 1 µg repair DNA fragment as described (Mans et al., 2015), followed by selection on a ura^-^ his^-^ plate. The correct genetic modification was verified by restriction analysis and by partial sequencing of a 1725bp DNA fragment (Chr XIV 656333-658057) containing the mutation, that was obtained by PCR amplification of genomic DNA using the primers listed (Table S2). Digestion of the PCR mixture with *Psp*1406I*/*AclI (ThermoFisher Scientific) according to the manufacturer’s instructions confirmed the loss of a restriction site. The purified PCR product was partially sequenced by Eurofins (Ebersberg, Germany) with the forward primer (Table S2) that starts at position 657469 of the Crick chain of chromosome XIV. All primers were obtained from IDT (Leuven, Belgium). Plasmids were removed by culturing the cells in liquid SD medium containing 20 μg/mL uracil and 20 μg/mL histidine for 7 days/passages, and verified by growth on the non-selective (ura^+^ his^+^) plate but not on selective (ura^-^ and his^-^) medium.

The loss of one copy of chromosome XV from a diploid *cho2opi3* strain was induced using a conditional centromere as described (Reid et al., 2008). Briefly, plasmid pCEN15-UG was digested with *NotI* (ThermoFisher Scientific) to liberate the integrating fragment containing the *CEN* locus of Chr XV interrupted by the *K. lactis URA3* gene and the *GAL1* promoter. The fragment was transformed into *co MATα*, generating *co MATα CEN15*::*pGal1-CEN15-Kl_URA3* by homologous recombination (Guthrie and Fink, 1991). Transformants containing the conditional centromere were verified by PCR using the primers listed in Table S2. *co MATα CEN15::pGal1-CEN15-Kl_URA3* was crossed with *co MAT****a*** and diploids were selected on lys^-^ met^-^ medium as above. The conditional centromere was destabilized by plating on YPGal, and subsequently a copy of chromosome XV was removed by counter-selection against the *URA3* gene by replica-plating on SD medium containing 1 mg/mL 5-fluoroorotic acid (5-FOA, ThermoFisher Scientific) from a 100 mg/mL stock in DMSO. Plasmids were transformed according to the high efficiency transformation protocol (Guthrie and Fink, 1991). When removing the *URA3* gene by 5-FOA, strain growth was rescued with 20 μg/mL uracil.

### DNA content analysis by FACS

Samples for Fluorescence-Activated Cell Sorting (FACS) were prepared according to published protocols (Haase and Reed, 2002; Pavelka et al., 2010). 1×10^7^ yeast cells (corresponding to 0.5 OD_600_ units) were harvested and fixed with 1 mL 70% cold ethanol while spinning at 4 °C overnight. After washing with 200 mM Tris-HCl pH 8.0, 2 mM EDTA and a second wash with 100 mM Tris-HCl pH 8.0, 2 mM EDTA, the cells in 1mL 100 mM Tris-HCl pH 8.0, 2 mM EDTA were sonicated for 10 min in an ice bath (Branson 3800), and then re-suspended in 1 mL RNAse solution (1 mg/mL RNAse A [Sigma-Aldrich] in 50 mM Tris-HCl pH 8.0, boiled for 15 min and allowed to cool to room temperature) for 6 h at 37 °C at 800 rpm (shaking incubator, Eppendorf). After RNAse A was removed by centrifugation, samples were collected and incubated in 500 μL of proteinase K solution (100 µg/mL proteinase K [Sigma-Aldrich] in 50 mM Tris-HCl pH 8.0, 2 mM CaCl_2_) at 55 °C at 800 rpm overnight. The samples were washed with 1 mL 50 mM Tris pH 8.0, 10 mM EDTA, followed by another washing step with 1 mL 50 mM Tris pH 8.0, 1 mM EDTA, and finally re-suspended in 1 mL 1 µM Sytox Green (ThermoFisher Scientific) in 50 mM Tris-HCl pH 7.5. After sonication for 10 min in an ice bath, samples were analyzed with a FACSCalibur flow cytometer (Becton Dickinson, NJ) with excitation at 488 nm, and sorted based on area (DNA-A) of the Sytox Green fluorescence signal, which was collected in the FL1 channel (530 ± 15 nm). Data were plotted as histograms showing the fluorescence distribution with FlowJo Software (Tree Star Inc., OR).

### Whole genome sequencing

Genomic DNA was isolated from 2 clones of each strain corresponding to 50 OD_600_ units with the Genomic DNA Buffer Set and Genomic-tip 100/G (QIAGEN) according to the manufacturer’s instructions. Briefly, cell pellets were re-suspended in Y1 buffer containing zymolyase and incubated at 30 °C at 200 rpm for 1 h. Spheroplasts were pelleted, re-suspended with G2 buffer containing 2 μg/mL RNAse A and 20 μg/mL proteinase K, and incubated in a water bath at 50 °C for 1 h. After centrifugation at 3000 g, the supernatant was collected and loaded on Genomic-tip 100/G which was equilibrated with buffer QBT. After washing twice with buffer QC, gDNA was eluted with 5 mL buffer QF and then precipitated with 3.5 mL cold isopropanol at -20 °C overnight. The gDNA was spun down at 10,000 g at 4 °C and washed with 100% cold ethanol. The gDNA dissolved in 50 μL 10 mM Tris-HCl pH 8.0, 1 mM EDTA was used for whole genome sequencing or qPCR analysis. The concentration of gDNA was determined by measuring absorbance using Nanodrop (ThermoFisher Scientific). Sequencing libraries with a mean insert size of 350 bp were constructed from 50 ng of genomic DNA, and sequenced with paired-end (2 × 250 bp) runs using an Illumina MiSeq instrument and V2 reagent kit to a minimal depth of 25× base coverage. Final library concentrations were measured using Qubit (ThermoFisher Scientific).

The alignment of sequencing reads was done using Burrows Wheeler Alignment (bwa 0.7.5a) (Li and Durbin, 2009) with settings ‘bwa mem -c 100 -M’ against the *S. cerevisiae* reference genome (S288C version R64-1-1). Mapped reads were sorted and duplicate reads were marked using the Sambamba v0.5.8 toolkit. Single nucleotide variants (SNVs), insertions and deletions (InDels) were called using GenomeAnalysisTK v3.4.46 HaplotypeCaller. Variants present in the parent *cho2opi3* strain were filtered out from all samples, as were variants between the two clones of one strain. SNVs were assigned a ‘FILTER’ flag using GenomeAnalysisTK v3.4.46 VariantFiltration using settings “--filterName SNP_LowQualityDepth --filterExpression ‘QD < 2.0’ --filterName SNP_MappingQuality -- filterExpression ‘MQ < 40.0’ --filterName SNP_StrandBias --filterExpression ‘FS > 60.0’ --filterName SNP_HaplotypeScoreHigh --filterExpression ‘HaplotypeScore > 13.0’ --filterName SNP_MQRankSumLow --filterExpression ‘MQRankSum < -12.5’ --filterName SNP_ReadPosRankSumLow --filterExpression ‘ReadPosRankSum < -8.0’ --clusterSize 3 - clusterWindowSize 35”. For further analysis only SNVs with the ‘FILTER’ flag set to ‘PASS’ were considered.

Copy number variation (CNV) was determined based on read depth using freec v7.2, the GC normalized ratios were used to construct karyotype’s using a 5Kb sliding window. All code used for these analyses is freely and openly available on github (github.com/UMCUGenetics/IAP/release/tag/v2.5.1).

### Karyotype analysis by qPCR

gDNA was isolated from cells cultured to OD_600_∼1 as described above, and diluted to 50 pg/μL. Primer pairs targeting intergenic regions proximal (≤25 kb) to the centromere within each arm of 5 chromosomes (chr01La, chr01Ra, chr04La, chr04Ra, chr06La, chr06Ra, chr09La, chr09Ra, chr15La, chr15Ra) designed as described (Pavelka et al., 2010), were obtained from IDT (Leuven, Belgium), and dissolved at 3.2 μM in water. The qPCR reactions were performed in 96-well plates using a ViiA™ 7 Real-Time PCR System (Applied Biosystems, ThermoFisher Scientific) in 20 μL reaction volumes. All reactions were set up in technical duplicates. Each reaction contained 10 μL Power SYBR Green PCR Master Mix (Applied Biosystems, ThermoFisher Scientific), 2.5 μL 3.2 μM forward primer, 2.5 μL 3.2 μM reverse primer, and 5 μL 50 pg/μL gDNA. The cycling conditions were 50 °C for 2 min, 95 °C for 10 min, followed by 40 cycles of 95 °C for 15 s and 60 °C for 1 min (Pavelka et al., 2010). Melting curves were recorded (15 s at 95 °C, 1 min at 60 °C, followed by a gradual increase to 95 °C in 15 s) to verify that no side products had been amplified. Ct values were determined using ViiA™7 Software.

Chromosome copy numbers were calculated as described previously using BY4742 as WT euploid reference strain (Livak and Schmittgen, 2001; Pavelka et al., 2010).

### Lipid extraction

Yeast total lipid extracts were prepared by the 2-phase lipid extraction method (Houtkooper et al., 2006), unless indicated otherwise. Briefly, cells corresponding to 200 OD_600_ units were lyophilized. The dry cell powder was resuspended in 6 mL chloroform: methanol (2:1, v/v) with 0.1 mL 0.1 M HCl, and sonicated for 20 min in a Branson B1200 bath sonicator (Bransonic Ultrasonics, Danbury, CT) containing ice water. Subsequently, 1 mL water was added to induce phase separation. After vigorous vortexing, cell debris was removed by centrifugation at 3000 g for 4 min. The organic phase was collected and residual lipid in the aqueous phase was extracted again with 3 mL chloroform: methanol (2:1, v/v). After washing with water, the combined organic phase was dried under a stream of nitrogen. The phospholipid content of lipid extracts was determined as described (Rouser et al., 1970) using KH_2_PO_4_ as a standard after destruction in 70% HClO_4_ at 180 °C for 2 h.

### Thin layer chromatography

*Separation of phospholipids by 2D-TLC* Total lipid extracts corresponding to 200 nmol phospholipid were applied on silicagel plates (Merck 1.05641), freshly impregnated with 2.4% (w/v) boric acid (de Kroon et al., 1997). The eluent for the first dimension contained chloroform: methanol: 25% ammonia (71:30:4, v/v/v). After drying under a flow of nitrogen for 30 min, the plate was run in the second dimension using chloroform: methanol: acetic acid (70:25:10, v/v/v) as eluent. The lipid spots were visualized by iodine staining. Spots were scraped off and phospholipid classes were quantified (Rouser et al., 1970).

*Separation of neutral lipids by 1D-TLC* Total lipid extracts corresponding to 10 nmol phospholipid were applied on a silicagel plate. The neutral lipids were separated using hexane: diethylether: acetic acid (35:15:1, v/v/v) as eluent (Schneiter and Daum, 2006). Ergosterol obtained from Sigma-Aldrich (E-6625), cholesterol ester, monoacylglycerol (MAG), diacylglycerol (DAG), triacylglycerol (TAG) and free fatty acid obtained from Nu-Chek-Prep (Elysian, MN) were used as standards. The lipid spots were visualized by MnCl_2_ charring.

### ^32^P-labeling

10 OD_600_ units of mid-log phase cells pre-cultured as above were harvested, washed and resuspended in 2.5 mL SD medium with or without 1 mM choline. After 30 min at 30 °C in a shaking incubator, [^32^P]orthophosphate (8500 Ci/mmol, Perkin Elmer) was added to 100 μCi/mL. The incubation was continued for 30 min and ended by adding 5% (w/v) TCA and putting the samples on ice. Cells were washed with water twice, homogenized by vortexing in the presence of glass beads for 3 times 1.5 min with intermittent cooling on ice. After adding HCl to 0.1 M, lipids were extracted (Bligh and Dyer, 1959). To determine the distribution of the lipid-incorporated [^32^P]-label over the phospholipid classes, the lipid extracts were analyzed by 2D-TLC as above. Radioactive spots were detected using a Typhoon FLA7000 PhosphorImager (GE Healthcare Life Sciences) and quantified by ImageQuant TL8.1 software.

### Gas chromatography

Total lipid extracts corresponding to 100 nmol of phospholipid phosphorus were transesterified in 3 mL methanol containing 2.5% (v/v) sulfuric acid at 70°C for 2.5 h. After cooling to room temperature, 2.5 mL water and 2.5 mL hexane were added. The hexane phase was collected and the aqueous phase was washed with another 2.5 mL hexane. After washing the pooled hexane phase at least three times with water to remove residual sulfuric acid, 100 µL isopropanol was added, and the samples were dried under nitrogen gas. 100 µL of hexane was added to the fatty acid methyl esters (FAME).

FAME were analyzed by Gas Chromatography-Flame Ionisation Detection (GC-FID) on a Trace GC Ultra (ThermoFisher Scientific) equipped with a biscyanopropyl polysiloxane column (Restek, Bellefonte PA) using nitrogen as carrier gas and a temperature gradient from 160 °C to 220 °C. Peak identification and calibration of the integrated signal intensities were performed using a FAME standard (63-B, Nu-Chek-Prep).

### Lipid extraction for mass spectrometry-based lipidomics

Cells were harvested at mid-log phase (OD_600_ ∼0.3-1) and washed twice with 150 mM ice cold ammonium bicarbonate (ABC) buffer. Cells corresponding to ∼10 OD_600_ units were resuspended in 0.5 mL ABC, and vortexed vigorously in the presence of 200 μL glass beads for 2 x 5 min at 4°C with intermittent cooling on ice for 3 min. The lysates were frozen directly in liquid nitrogen and stored at -80 °C until further processing. Mass spectrometry-based lipid analysis was performed by Lipotype GmbH (Dresden, Germany) as described (Ejsing et al., 2009; Klose et al., 2012). Lipids were extracted using a two-step chloroform/methanol procedure (Ejsing et al., 2009). Samples were spiked with internal lipid standard mixture containing: CDP-DAG 17:0/18:1, ceramide 18:1;2/17:0 (Cer), diacylglycerol 17:0/17:0 (DAG), lysophosphatidate 17:0 (LPA), lyso-phosphatidylcholine 12:0 (LPC), lysophosphatidylethanolamine 17:1 (LPE), lyso-phosphatidylinositol 17:1 (LPI), lysophosphatidylserine 17:1 (LPS), phosphatidate 17:0/14:1 (PA), phosphatidylcholine 17:0/14:1 (PC), phosphatidylethanolamine 17:0/14:1 (PE), phosphatidylglycerol 17:0/14:1 (PG), phosphatidylinositol 17:0/14:1 (PI), phosphatidylserine 17:0/14:1 (PS), ergosterol ester 13:0 (EE), triacylglycerol 17:0/17:0/17:0 (TAG), stigmastatrienol, inositolphosphorylceramide 44:0;2 (IPC), mannosylinositolphosphorylceramide 44:0;2 (MIPC) and mannosyl-di-(inositolphosphoryl)ceramide 44:0;2 (M(IP)_2_C). After extraction, the organic phase was transferred to an infusion plate and dried in a speed vacuum concentrator. 1^st^ step dry extract was re-suspended in 7.5 mM ammonium acetate in chloroform/methanol/propanol (1:2:4, v/v/v) and 2^nd^ step dry extract in 33% ethanol solution of methylamine in chloroform/methanol (0.003:5:1, v/v/v). All liquid handling steps were performed using Hamilton Robotics STARlet robotic platform with the Anti Droplet Control feature for organic solvents pipetting.

### MS data acquisition

Samples were analyzed by direct infusion on a QExactive mass spectrometer (Thermo Scientific) equipped with a TriVersa NanoMate ion source (Advion Biosciences). Samples were analyzed in both positive and negative ion modes with a resolution of R_m/z=200_=280,000 for MS and R_m/z=200_=17,500 for MSMS experiments, in a single acquisition. MSMS was triggered by an inclusion list encompassing corresponding MS mass ranges scanned in 1 Da increments (Surma et al., 2015). Both MS and MSMS data were combined to monitor EE, DAG and TAG ions as ammonium adducts; PC as an acetate adduct; and CL, PA, PE, PG, PI and PS as deprotonated anions. The absence of PC in *cho2opi3* suppressor strains was confirmed in positive ion mode by MSMS scan for the phosphocholine headgroup fragment. MS only was used to monitor LPA, LPE, LPI, LPS, IPC, MIPC, M(IP)_2_C as deprotonated anions; Cer and LPC as acetate adducts and ergosterol as protonated ion of an acetylated derivative (Liebisch et al., 2006).

### Data analysis and post-processing

Data were analyzed with in-house developed lipid identification software based on LipidXplorer (Herzog et al., 2011, 2012). Data post-processing and normalization were performed using an in-house developed data management system. Only lipid identifications with a signal-to-noise ratio >5, and a signal intensity 5-fold higher than in corresponding blank samples were considered for further data analysis. Since the TAG species cannot be unambiguously assigned because of the presence of 3 FAs (is combinatorically not possible), the acyl chain distribution of the TAG fraction was quantitated as follows. TAG precursors were fragmented and the FAs released identified. The profile of FAs determined for each TAG precursor was normalized to the abundance of the precursor, yielding pmol values for each FA, which were summed up to obtain the FA profile for TAG. In quantifying the lipidomics data, molecular species containing acyl chains with an odd number of C atoms or a number of C atoms larger than 18 that constituted less than 3% and 0.5% of total, respectively, were left out.

### Electron microscopy

The preparation of samples for EM analysis was performed as described previously (Griffith et al., 2008; Mari et al., 2014). Briefly, after harvest the cells were rapidly mixed with an equal volume of double strength fixative [4% (w/v) paraformaldehyde (PFA, Sigma-Aldrich), 0.4% (v/v) glutaraldehyde (GA, Polyscience Inc) in 0.1 M Phem [20 mM PIPES, 50 mM HEPES, pH 6.9, 20 mM EGTA, 4 mM MgCl_2_] pH 6.9, and incubated for 20 min at room temperature on a roller bank. The mixture fixative-media was replaced by 2% PFA, 0.2% GA in 0.1 M Phem pH 6.9 for 2 h at RT followed by an overnight incubation at 4°C. Fixative was removed by centrifugation and cells were embedded in 12% gelatin. Blocks of 1 mm^3^ were obtained and incubated in 2.3 M sucrose overnight at 4°C before being mounted on pins. Cells sections were obtained using a LEICA cryo-EM UC7. Membrane contrast was performed as described previously (Griffith et al., 2008; Mari et al., 2014). Thin sections (50 nm) were viewed in an electron microscope (1200 EX; JEOL).

The 2D projection images (Tiff format) of non-overlapping regions in the cryosection were imported into the IMOD software package. Cell area and lipid droplet area were determined in at least 15 2D projection images by counting the total number of pixels covering yeast cells and the total number of pixels covering the lipid droplets. Lipid droplet content is expressed as the area occupied by lipid droplets determined as a percentage of total cell area and was calculated using Excel. Significance was determined with Student’s t-test.

### mRNA profiling

Wild-type BY4742, *co* S#3, S#4 and S#5 were precultured in SD medium (C^-^) to mid-log phase (OD_600_ ∼1.0), and transferred to 15 mL of the corresponding medium at OD_600_ of 0.05. The *co MATα* parent strain was precultured to OD_600_ ∼1.0 in SD medium containing 1 mM choline; cells were collected by filtration, washed with choline-free SD medium (30°C), and used to inoculate 15 mL fresh SD C^-^ medium to an OD_600_ of 0.1 (Boumann et al., 2006). All strains were cultured at 30°C in biological replicate and harvested in early mid-log phase (OD_600_ of 0.55-0.65) by centrifugation at room temperature for 3 min. Time from removing culture from incubator until freezing pellet is maximally 5 min.

Total RNA was isolated by phenol extraction and purified as described (Kemmeren et al., 2014). All subsequent procedures in expression-profiling including RNA amplification, cRNA labeling, microarray hybridization, quality control, and data normalization were carried out as described previously (Kemmeren et al., 2014). Two channel microarrays were used. RNA isolated from WT was used in this common reference design, in one of the channels for each hybridization. Two independent cultures were hybridized on two separate microarrays. For the first hybridization the Cy5 (red) labeled cRNA from the mutant was hybridized together with the Cy3 (green) labeled cRNA from the common reference. For the replicate hybridization from the independent cultures, the labels were swapped. The reported fold change is the average of the four replicate mutant profiles versus the average of the WT controls. Genes were considered significantly changed when the fold-change (FC) was > 1.7 and the p value < 0.01. P values were obtained from the limma R package version 2.12.0 (Smyth, 2005) after Benjamini-Hochberg FDR correction. Hierarchical clustering of genes subject to significant expression changes was by average linkage (cosine correlation). Functional enrichment was by a hypergeometric test on Gene Ontology Biological Process (GO-BP; p<0.01, Bonferroni corrected). Enriched GO terms were summarized by the REVIGO software using a cutoff value C of 0.5 (Supek et al., 2011).

### RT-qPCR

Diploid yeast strain BY4743, *co* diploid, *co* S(2n-1) and *co* S#2 were precultured in SD C^+^I^+^ to log phase (OD_600_ ∼0.45-1.2). After washing with pre-warmed SD C^-^I^-^ (30°C) by filtration, cells were rapidly transferred to C^+/-^I^+/-^ and cultured for 24 h. Total RNA was isolated from 20 OD_600_ units after cell disruption by rapid agitation in the presence of glass beads and lysis buffer, using the RNeasy™ Mini Kit and RNase-free DNase Set according to the manufacturer’s instructions (QIAGEN). RNA quality and quantity were checked on a 1% agarose gel and with a NanoDrop spectrophotometer (ND-1000, ThermoFisher Scientific), respectively.

1 µg RNA was converted to cDNA according to the first strand cDNA synthesis protocol of Invitrogen™ (ThermoFisher Scientific) using SuperScript™ III reverse transcriptase. 1 uL oligo(dT)12-18 Primer, 1 uL 10 mM dNTP and 1ug RNA were mixed in a PCR tube, and water was added to adjust the total volume to 14 uL. Mixture was heated at 65°C for 5 min and incubated on ice for 1 min. 4 uL 5x First Strand Buffer, 1 uL 0.1 M DTT and 1 uL SuperScriptTM III Reverse Transcriptase was added into the mixture gently, followed by an incubation at 50°C for 60 min. The reaction was inactivated by heating at 70°C for 15 min, and 80 uL MQ water was added.

qPCR was performed in technical duplicate as described above in 96 wells plates with 20 uL reactions containing 10 uL TaqManTM Universal PCR Master Mix, no AmpEraseTM UNG (Applied Biosystems, ThermoFisher Scientific), 1 uL TaqMan™ probes and primers (*ACT1, ACC1, POX1, KAR2, INO1*, Reagents and Tools Table), 4 uL MQ water and 5 uL cDNA (from 50 ng RNA). Non-template control (cDNA) and non-reaction control were routinely performed. The data was analyzed according to the ΔΔCt method (Livak and Schmittgen, 2001), normalized to the control gene *ACT1*, and expressed relative to the corresponding strain cultured in C^+^I^+^.

### Quantification and statistical analysis

For each experiment the number of biological replicates is indicated in the corresponding figure legend and/or methods section. GraphPad Prism 8 was used to determine means, standard deviations and statistical significance in the column graphs, with p-values determined by an unpaired two-tailed t-test, with * p < 0.05, ** p < 0.01, *** p < 0.001, **** p < 0.0001.

### Data availability

The raw sequencing data are available as BAM files at the European Nucleotide Archive (ENA) through accession number PRJEB40705 (http://www.ebi.ac.uk/ena/data/view/PRJEB40705).

The transcript profiling data reported in this publication are accessible through GEO Series accession number GSE75725 (http://www.ncbi.nlm.nih.gov/geo/query/acc.cgi?acc=GSE75725).

*The following secure token has been created to allow review of record GSE75725 while it remains in private status:* gzmlyacadryjtuf. *(Delete this line from the final preprint version of the paper!!)*

## ACKNOWLEDGMENTS

We thank Wouter Beugelink, Robin Hoogebeen, and Linda Markus for help with strain construction, Ivo Renkens, Ies Nijman, and Joep de Ligt for WGS assistance, Ger Arkesteijn for FACS assistance, Jingchao Wu and Yifei Lang for assistance with gene editing by CRISPR/Cas9, Robert Reid and Klaus Natter for plasmids, Marc Stadler (Helmholtz Centre for Infection Research, Braunschweig, DE) for a kind gift of Soraphen A, Christoph Thiele for providing ergosterol ester C13:0, Philip Lijnzaad, Henk van den Toorn, and Bas van Breukelen for assistance with the analysis of the mRNA profiling data, and Sepp Kohlwein, Klaus Natter, and Harald Hofbauer for valuable discussions. We are indebted to Antoinette Killian and Fulvio Reggiori for critically reading the manuscript.

We thank Utrecht Sequencing Facility for providing sequencing service and data. Utrecht Sequencing Facility is subsidized by the University Medical Center Utrecht, Hubrecht Institute and Utrecht University. We thank UMC Utrecht Bioinformatics Expertise Core for data analysis and data handling. The UMC Utrecht Bioinformatics Expertise Core is subsidized by the University Medical Center Utrecht, Center for Molecular Medicine. This research was supported by the Division of Chemical Sciences in the Netherlands, with financial aid from The Netherlands Organization for Scientific Research (711.017.010, XB) and by the China Scholarship Council (grant no. 201204910146, XB).

## AUTHOR CONTRIBUTIONS

Conceptualization: X.B., A.I.P.M.d.K.; Funding acquisition: X.B., A.I.P.M.d.K.; Investigation: X.B., M.C.K., M.J.A.G.K., A.H., A.W., S.C., M.F.R., W.J.C.G., M.A.S., M.M.; Methodology: X.B., M.J.A.G.K., W.J.C.G., M.A.S., M.M., F.C.P.H., C.K., A.I.P.M.d.K.; Supervision: X.B., A.I.P.M.d.K., Vi*s*ualization: X.B., M.M., A.I.P.M.d.K., Writing-original draft: X.B., A.I.P.M.d.K.; Writing-review & editing: all authors.

## Supplemental information

Supplemental Information includes seven figures, two tables and three datasets.

Data S1, Nucleotide variation identified by WGS in the *cho2opi3* suppressors S#2, S#3, S#4 and S#5 *versus* the haploid parent strain

Data S2, Lipidome analyses of WT, *cho2opi3* and derived suppressor strains cultured in the absence and/or presence of choline

Data S3, Transcript profiling data of the *cho2opi3* parent strain and evolved suppressors S#3, S#4, and S#5

## REAGENTS AND TOOLS TABLE

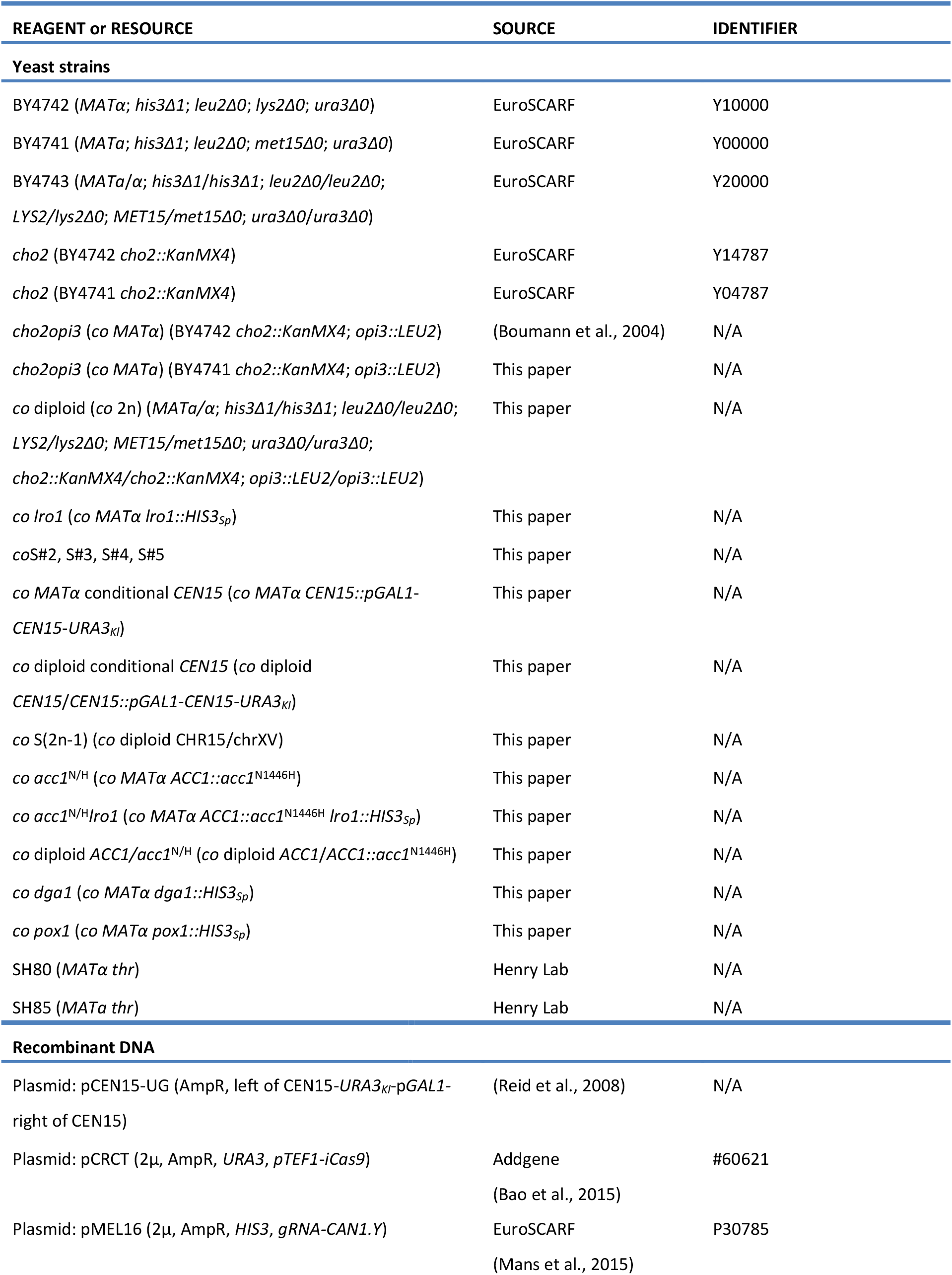
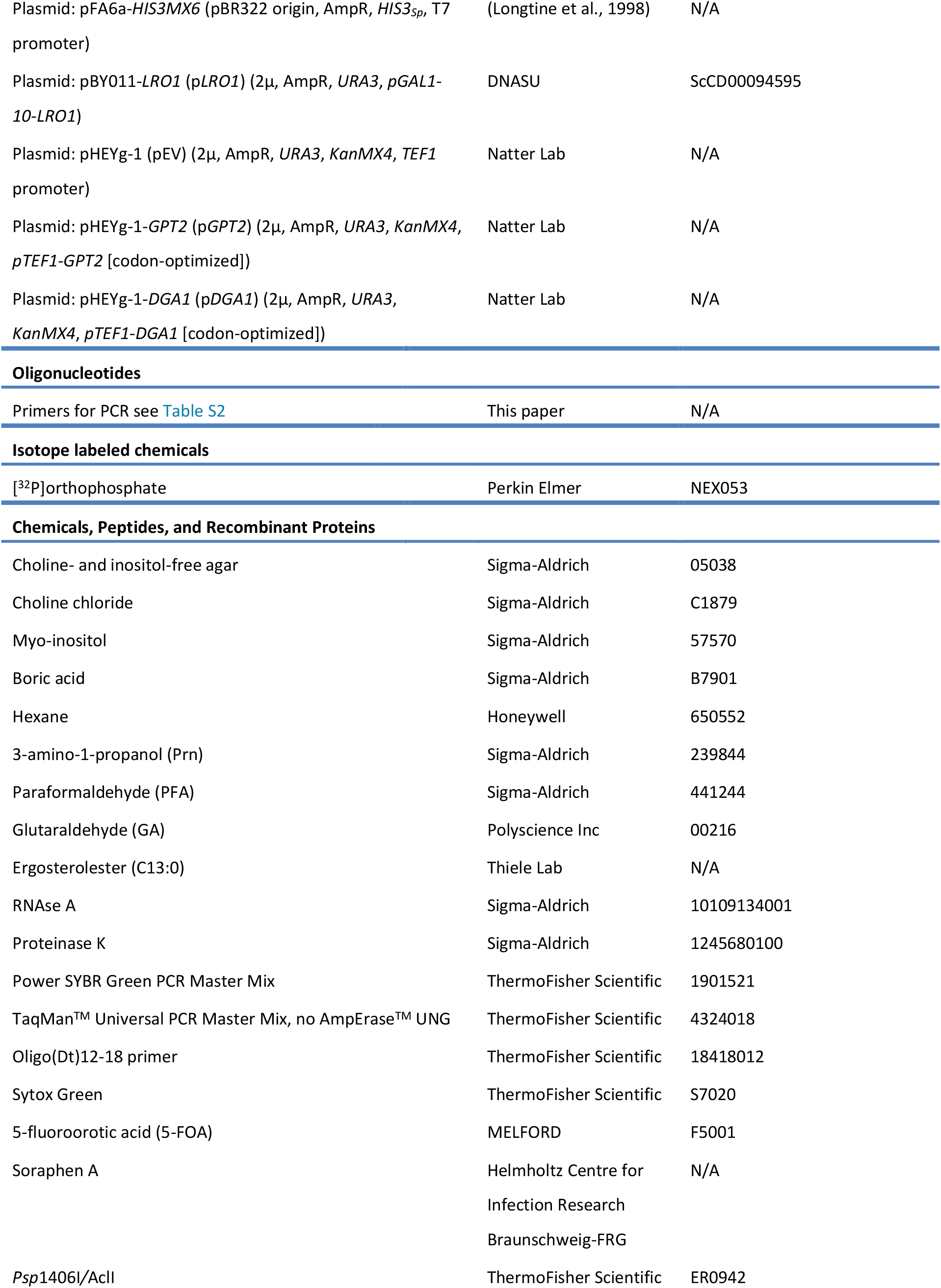
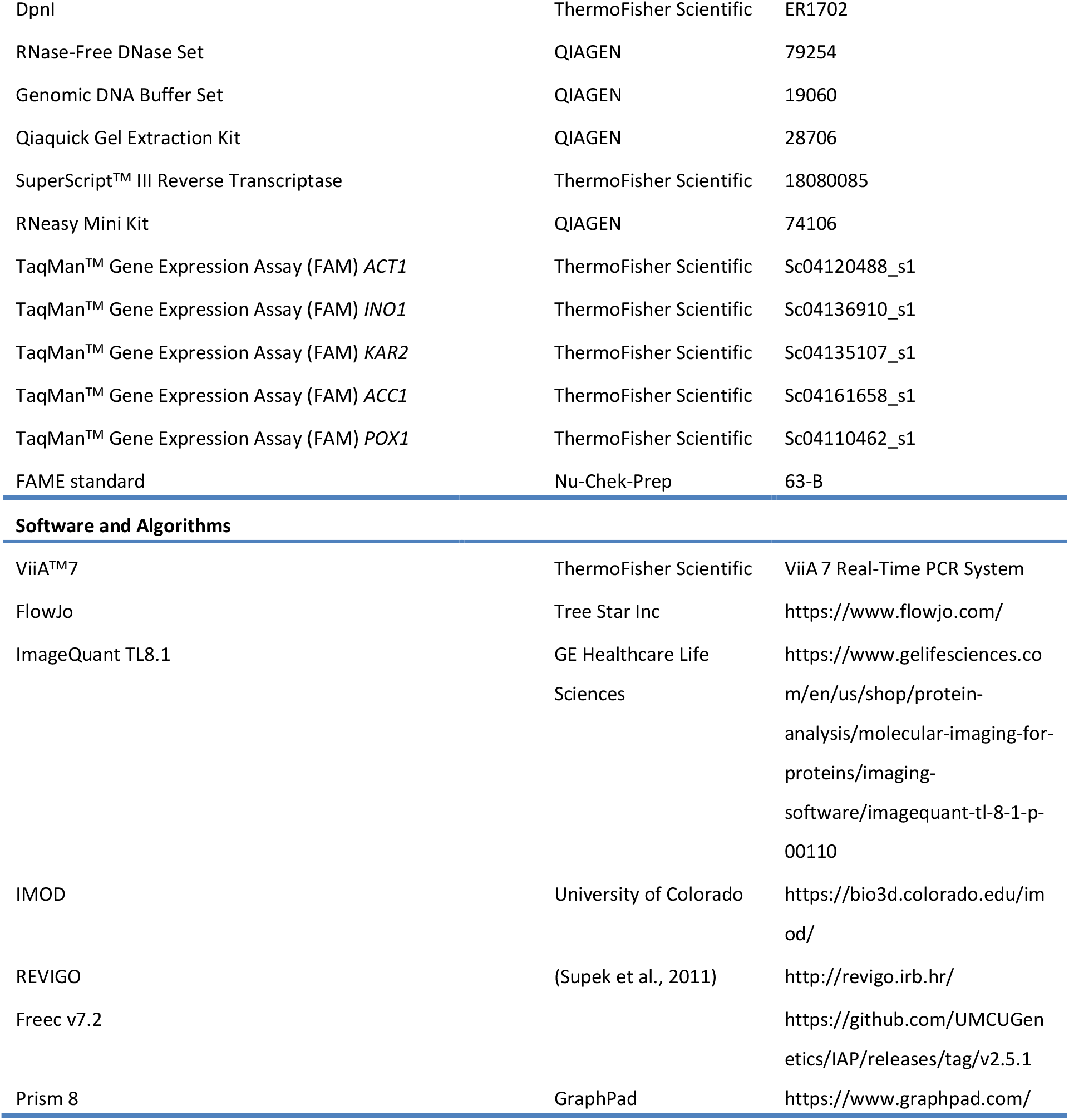

