## Supplemental Information for "Acyl chain shortening induced by inhibition of acetyl-CoA carboxylase renders phosphatidylcholine redundant"

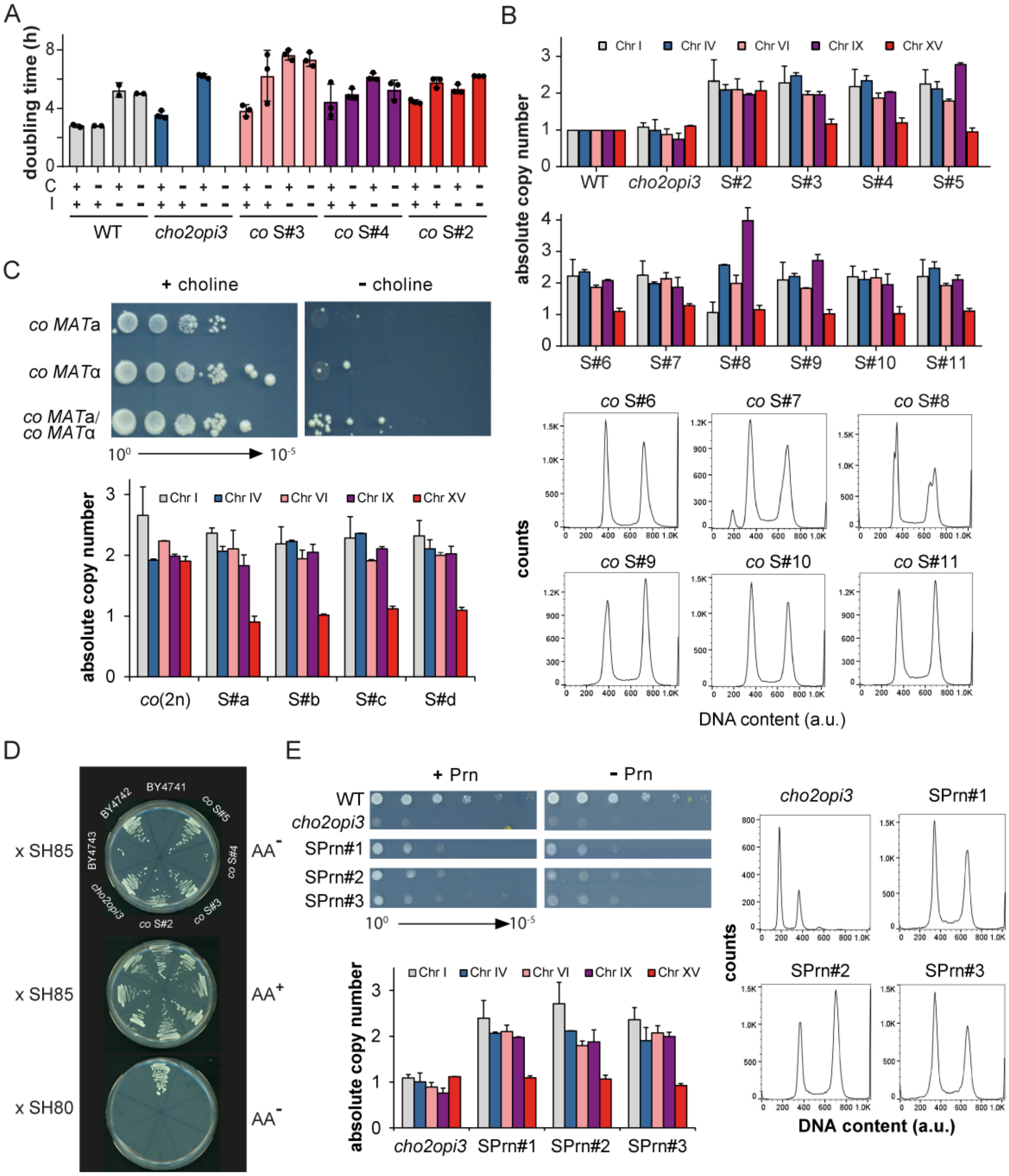


Figure S1

**Figure S1. Characterization of Evolved *cho2opi3* Suppressors, Related to Figure 1**

(A) Doubling times of wild type, *cho2opi3*, and *co* S#3, S#4 and S#2, cultured in SD medium supplemented with 1 mM choline and/or 75 $\mu$M inositol as indicated. The error bars represent SD (wild type, n=2; other strains, n=3).

(B) 9 out of 10 independent *cho2opi3* suppressor strains exhibit monosomy of chr XV. Copy numbers of chr I, IV, VI, IX and XV were derived from qPCR and FACS analysis of cellular DNA content (lower panel and Fig. 1E). The error bars represent the variation between assays using primers complementary to non-coding regions on the left and right arm of each chr, respectively.

(C) Serial dilution experiment ($10^0$ - $10^{-5}$) comparing haploid *co MAT**a*** and *co MAT*$\alpha$ to *co* diploid on SD $C^{+/-}$ after incubation for 14 d at 30°C, and absolute copy numbers of chr I, IV, VI, IX and XV in a *co* diploid and 4 derived suppressor strains as determined by qPCR and FACS.

(D) The *cho2opi3* suppressor strains retain the $\alpha$-mating type. Suppressor and control strains were mated with SH85 (*MAT**a***) and SH80 (*MAT*$\alpha$) as indicated, and then streaked on SD $C^-$ plates without amino acids ($AA^-$). The amino acid containing plate ($AA^+$) serves as control. The reduced mating efficiency of *co*S#4 may be due to haplo-insufficiency originating from the loss of one copy of the *MAT* locus on chr III. The ability of BY4743 (*MAT**a***/*MAT*$\alpha$) to mate with SH85 is attributed to loss of heterozygosity by homologous recombination (Harari et al., 2018).

(E) Characterization of 3 *cho2opi3* suppressor clones generated on SD $C^-I^-$ plates supplemented with 1 mM propanolamine (Prn). Growth phenotype on SD with 1 mM Prn or without supplement after 5 d at 30°C, DNA content by FACS, and absolute copy numbers of chr I, IV, VI, IX and XV in *co* SPrn#1, #2 and #3 compared to wild type and the haploid *co* parent. The error bars represent the variation between qPCR assays using primers complementary to non-coding regions on the left and right arm of each chr, respectively.

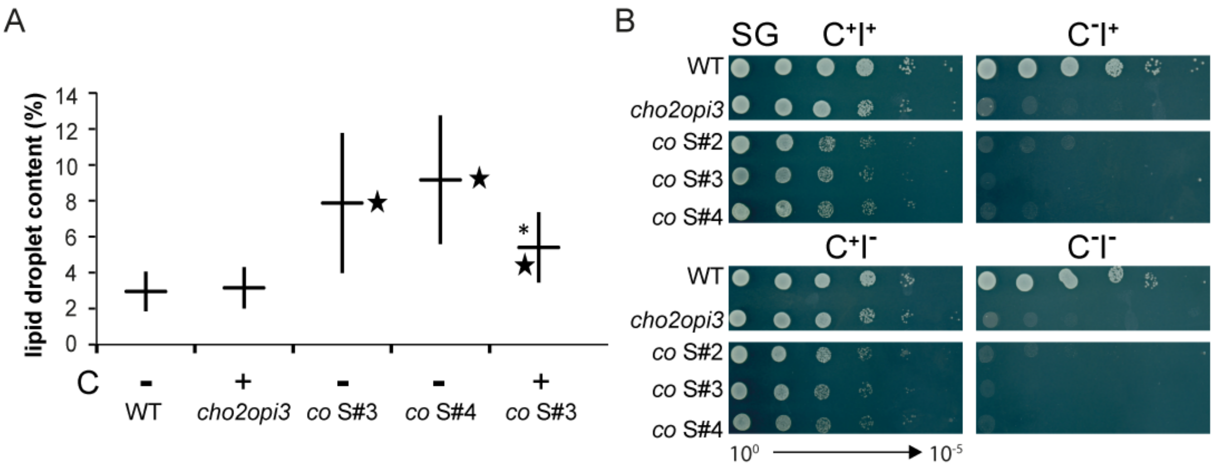


Figure S2

**Figure S2. Phenotypes of 2n-1 Evolved *cho2opi3* Suppressors, Related to Figure 1**

(A) Quantification of lipid droplet size in wild type, *cho2opi3*, *co*S#3 and S#4 after culture to mid-log phase in SD $C^{+/-}$ as indicated. Lipid droplet relative size was determined as a percentage of total cell area in 15 2D projection images. Details are described in STAR*Methods; error bars represent the SD (n=15); ★ represents P-value < 0.0002 *vs*. wild type; * represents P-value <0.02 *vs*. S#3 without choline.

(B) *cho2opi3* suppressors do not grow on the non-fermentable carbon source glycerol. Ten-fold serial dilutions of 1 $OD_{600}$ unit/mL of the strains indicated were spotted on SG $C^{+/-}$ $I^{+/-}$ plates, and incubated at 30°C for 10 d.

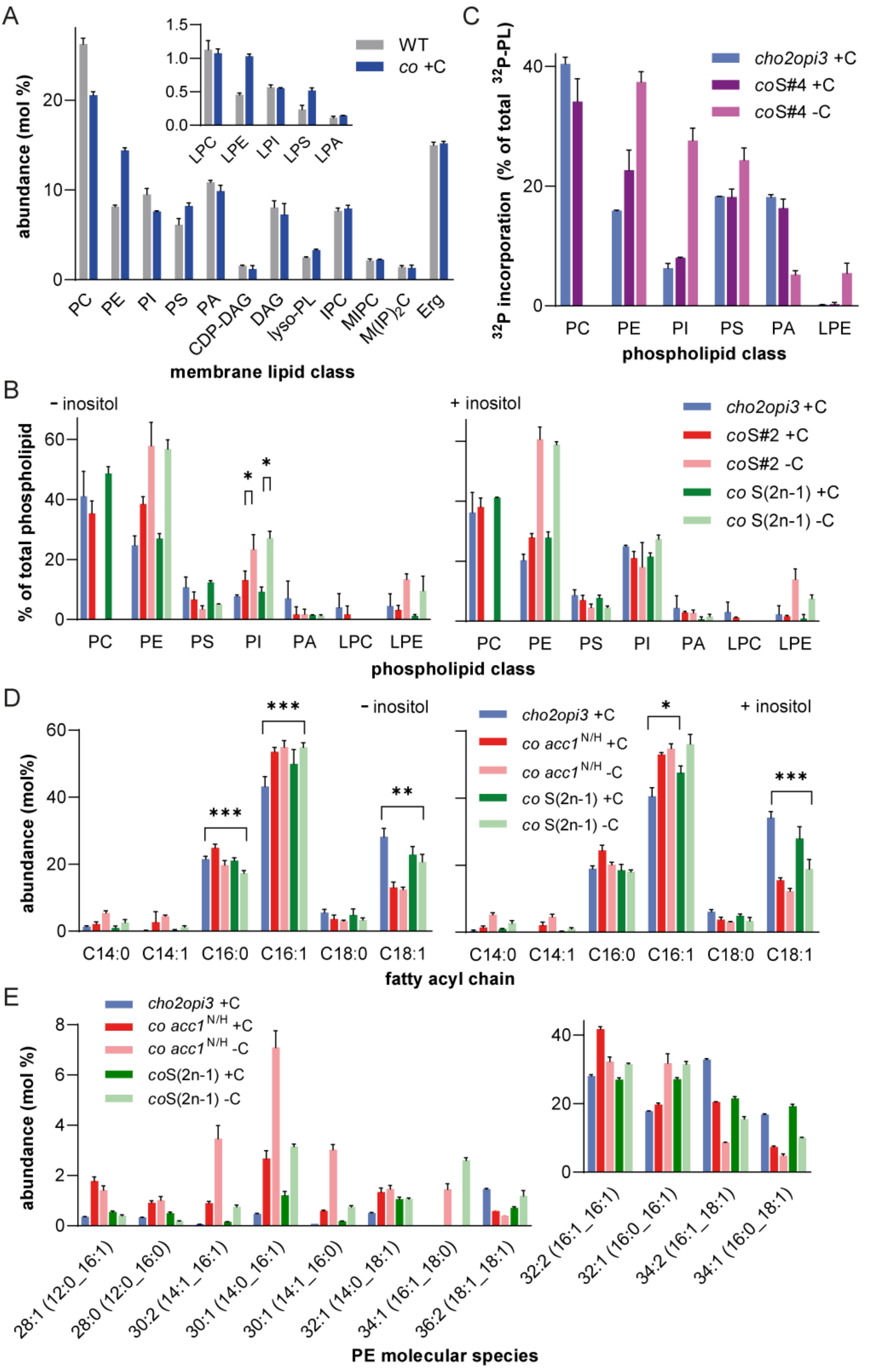
A
abundance (mol %)
WT
co +C
PC PE PI PS PA CDP-DAG DAG lyso-PL IPC MIPC M(IP)2C Erg
LPC LPE LPI LPS LPA
membrane lipid class
C
32P incorporation (% of total 32P-PL)
cho2opi3 +C
coS#4 +C
coS#4 -C
PC PE PI PS PA LPE
phospholipid class
B
- inositol
+ inositol
% of total phospholipid
cho2opi3 +C
coS#2 +C
coS#2 -C
co S(2n-1) +C
co S(2n-1) -C
PC PE PS PI PA LPC LPE
phospholipid class
D
- inositol
+ inositol
abundance (mol%)
cho2opi3 +C
co acc1 N/H +C
co acc1 N/H -C
co S(2n-1) +C
co S(2n-1) -C
C14:0 C14:1 C16:0 C16:1 C18:0 C18:1
fatty acyl chain
E
abundance (mol %)
cho2opi3 +C
co acc1 N/H +C
co acc1 N/H -C
coS(2n-1) +C
coS(2n-1) -C
28:1 (12:0_16:1)
28:0 (12:0_16:0)
30:2 (14:1_16:1)
30:1 (14:0_16:1)
30:1 (14:1_16:0)
32:1 (14:0_18:1)
34:1 (16:1_18:0)
36:2 (18:1_18:1)
32:2 (16:1_16:1)
32:1 (16:0_16:1)
34:2 (16:1_18:1)
34:1 (16:0_18:1)
PE molecular species

Figure S3

**Figure S3. Characterization of the Lipidome of PC-free *cho2opi3* Suppressors, Related to Figure 3**

(A) Membrane lipid class composition of *cho2opi3* cells cultured in SD $C^+$ compared to WT cultured in SD $C^-$, as analyzed by MS (mean ±SD, n=3). The inset shows the distribution of the separate lyso-phospholipids (lyso-PL).

(B) Phospholipid class composition of the indicated strains cultured to mid-log phase in SD $C^{+/-}I^{+/-}$ as indicated. Total lipid extracts were separated by TLC and phospholipid classes quantified by phosphate content (S#2, mean ±SD, n=3; *cho2opi3* and *co* S(2n-1), mean ±SD, n=2).

(C) Phospholipid synthesis in *cho2opi3* and derived *co* S#4 cultured in SD $C^{+/-}$ as indicated. Mid-log phase cells were pulse-labeled with [$^{32}$P]-orthophosphate for 30 min as detailed in the experimental section. Total lipid extracts were analyzed by 2D-TLC followed by phosphor imaging. The label incorporated per phospholipid class has been depicted as percentage of the total label incorporated in the phospholipid fraction (mean ±SD, n=2) for those classes that contain at least 2% of the label in one of the strains/conditions.

(D) Fatty acyl chain composition of total lipid extracts of the indicated strains cultured to mid-log phase in SD $C^{+/-}I^{+/-}$ as indicated, analyzed by GC (mean percentages of total ±SD, n=3 for +C, n=4 for -C).

(E) Molecular species profile of PE showing individual acyl chain combinations that contribute at least 1% of total PE with the inset showing the most abundant species, of the indicated strains cultured to mid-log phase in in SD $C^{+/-}$, as determined by MS.

* $p < 0.05$, ** $p < 0.01$, *** $p < 0.001$, unpaired two-tailed t-test of the indicated bar compared to the *cho2opi3* parent unless indicated otherwise.

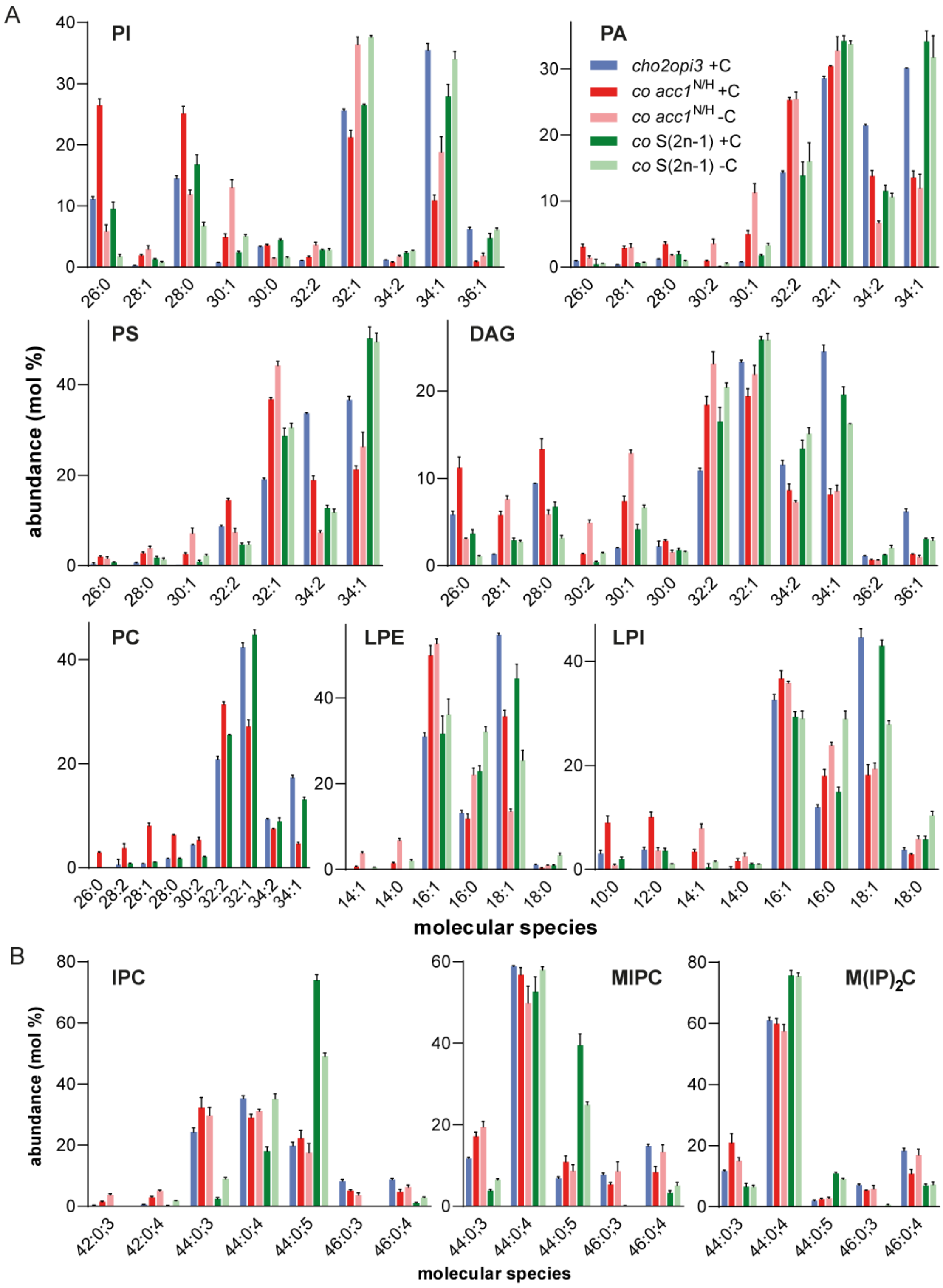


Figure S4

**Figure S4. Molecular Species Profiles of Membrane Lipids, Related to Figure 3**

Molecular species profiles (mol % of class) of (A) glycerolipid classes and (B) sphingolipid classes (sum of carbon atoms in the long chain base and fatty acyl chain : sum of double bonds in the long chain base and fatty acyl chain ; sum of hydroxyl groups in the long chain base and fatty acyl chain) in the strains indicated, cultured in in SD $C^{+/-}$. Molecular species contributing at least 2% of a class are depicted (± SD, n=3).

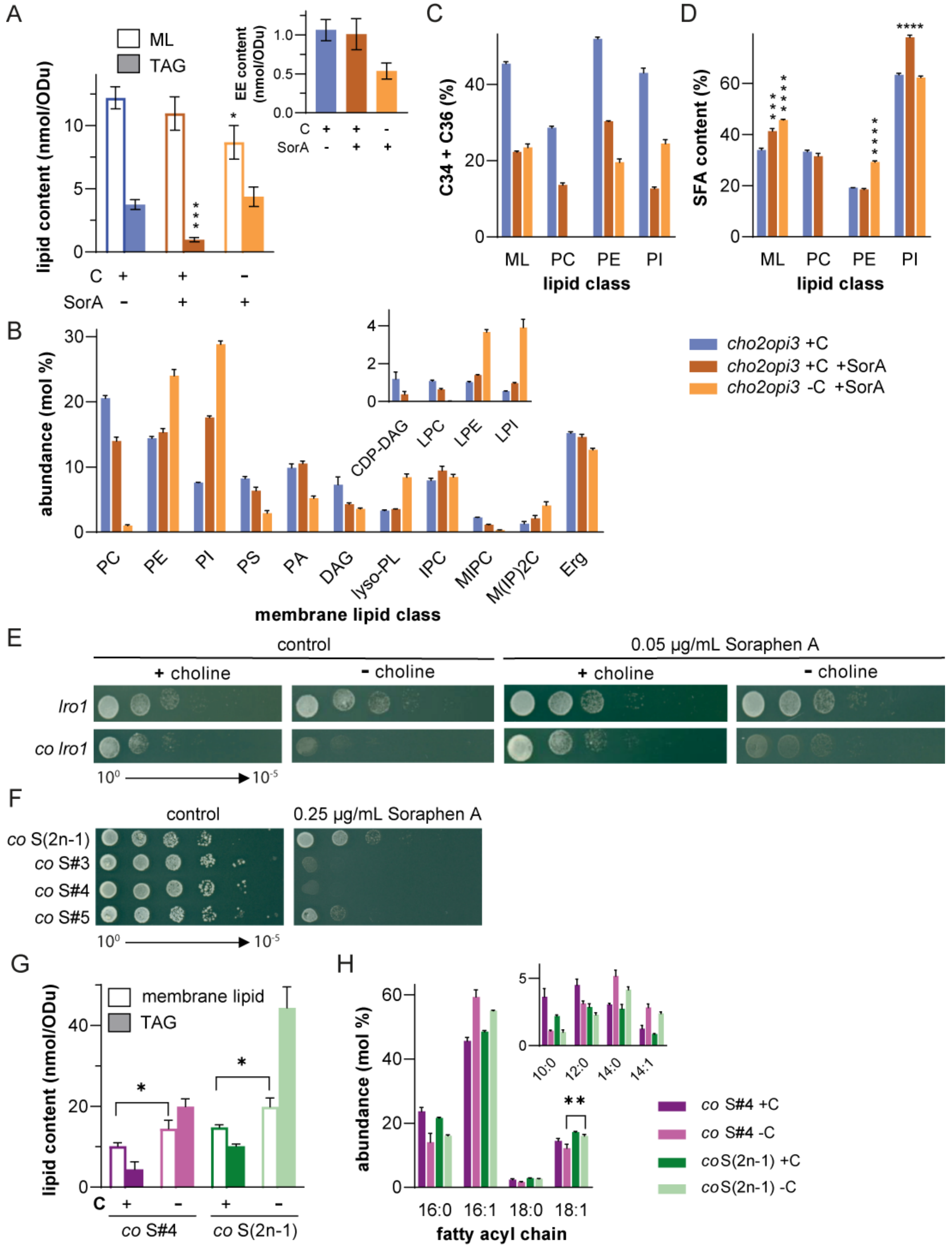
A
ML
TAG
lipid content (nmol/ODu)
C + + -
SorA - + +
EE content (nmol/ODu)
C
lipid class
C34 + C36 (%)
ML PC PE PI
D
SFA content (%)
B
abundance (mol %)
CDP-DAG LPC LPE LPI
cho2opi3 +C
cho2opi3 +C +SorA
cho2opi3 -C +SorA
PC PE PI PS PA DAG lyso-PL IPC MIPC M(IP)2C Erg
membrane lipid class
E
control
0.05 µg/mL Soraphen A
+ choline
- choline
lro1
co lro1
F
0.25 µg/mL Soraphen A
co S(2n-1)
co S#3
co S#4
co S#5
G
membrane lipid
co S#4
co S(2n-1)
H
abundance (mol %)
10:0 12:0 14:0 14:1
co S#4 +C
co S#4 -C
coS(2n-1) +C
coS(2n-1) -C
16:0 16:1 18:0 18:1
fatty acyl chain

Figure S5

**Figure S5. Effects of SorA on the Lipidome of *cho2opi3*, Related to Figure 5**

(A) Membrane lipid and TAG content, and EE content (inset) per $OD_{600}$ unit;

(B) membrane lipid class composition of lipid classes contributing at least 1% of total membrane lipids, with the inset showing CDP-DAG and the separate lyso-phospholipids (lyso-PL);

(C) percentage of molecular species containing more than 32 carbon atoms in both acyl chains (C34+C36);

(D) percentage of saturated acyl chains (SFA) in the membrane glycerolipid fraction (ML), and the major membrane lipid classes, of *cho2opi3* cultured in in SD $C^{+/-}$ and in the presence or absence of 0.05 µg/mL SorA, as indicated. Data in A-D were obtained by mass spectrometry and are presented as mean ±SD (n=3); * $p < 0.05$, *** $p < 0.001$, **** $p < 0.0001$, unpaired two-tailed t-test of the indicated bar compared to *cho2opi3* with choline.

(E) Growth phenotype of the single *lro1* mutant and *cho2opi3lro1* on SD $C^{+/-}$ with and without 0.05 ug/mL SorA as indicated at 30 °C for 3 d.

(F) Growth of the suppressor strains indicated in the presence and absence of 0.25 µg/ml SorA on $C^{+}I^{+}$ at 30 °C for 6 d. Control plates contain 0.02% (v/v) ethanol.

(G) Membrane lipid and TAG content per $OD_{600}$ unit of *co* S#4 compared to *co* S(2n-1) after culture to mid-log phase in SD $C^{+/-}$.

(H) Fatty acyl chain composition of total lipid extracts of *co* S#4 compared to *co* S(2n-1) cultured under the conditions indicated. Acyl chains that contribute at least 1% of total are depicted. Data in G-H were obtained by mass spectrometry and are presented as mean ±SD (n=3); * $p < 0.05$, p < 0.01, unpaired two-tailed t-test comparing the indicated bars.

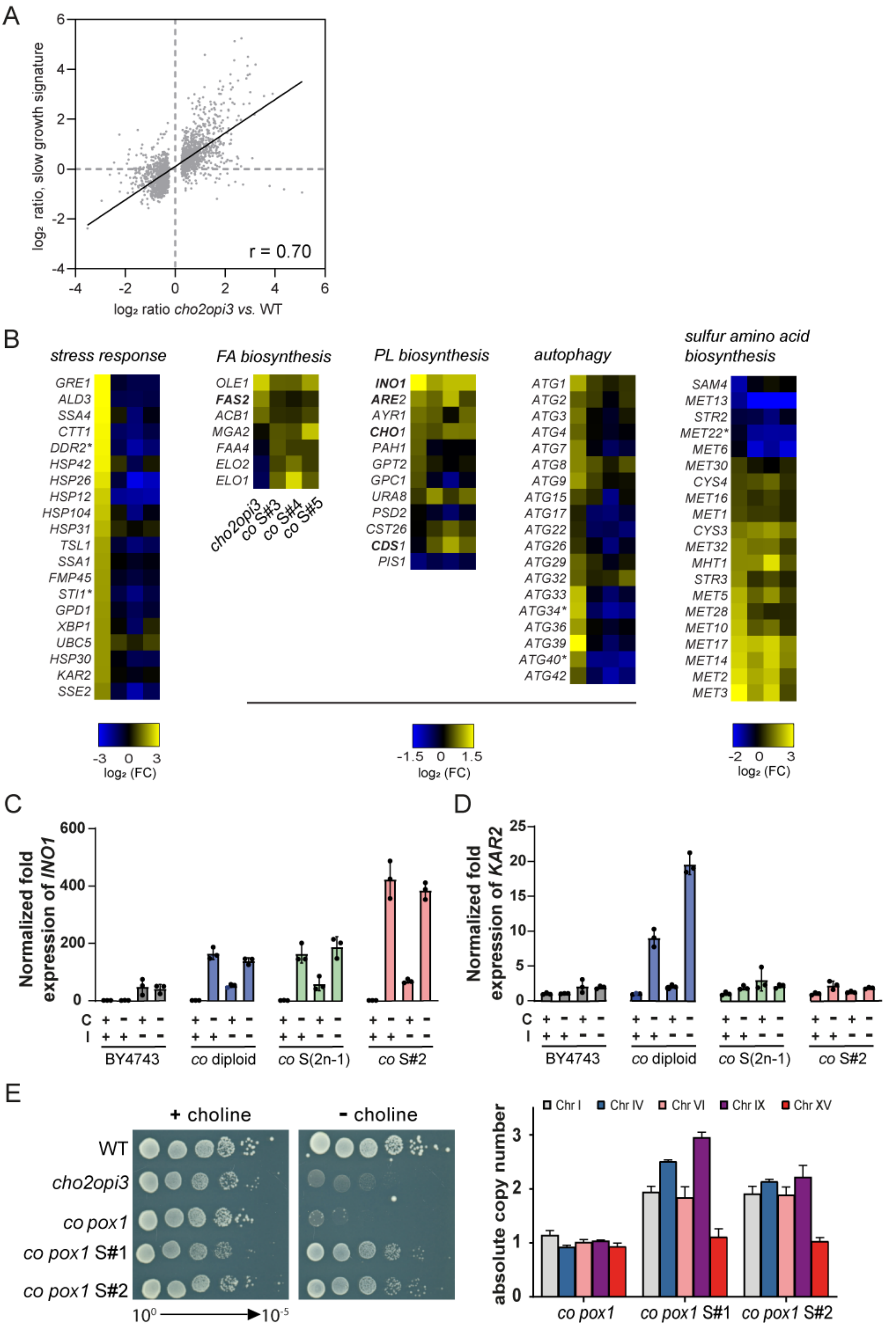

A
log₂ ratio, slow growth signature
log₂ ratio cho2opi3 vs. WT
r = 0.70
B
stress response
FA biosynthesis
PL biosynthesis
autophagy
sulfur amino acid biosynthesis
GRE1 ALD3 SSA4 CTT1 DDR2* HSP42 HSP26 HSP12 HSP104 HSP31 TSL1 SSA1 FMP45 STI1* GPD1 XBP1 UBC5 HSP30 KAR2 SSE2
OLE1 FAS2 ACB1 MGA2 FAA4 ELO2 ELO1
cho2opi3 co S#3 co S#4 co S#5
INO1 ARE2 AYR1 CHO1 PAH1 GPT2 GPC1 URA8 PSD2 CST26 CDS1 PIS1
ATG1 ATG2 ATG3 ATG4 ATG7 ATG8 ATG9 ATG15 ATG17 ATG22 ATG26 ATG29 ATG32 ATG33 ATG34* ATG36 ATG39 ATG40* ATG42
SAM4 MET13 STR2 MET22* MET6 MET30 CYS4 MET16 MET1 CYS3 MET32 MHT1 STR3 MET5 MET28 MET10 MET17 MET14 MET2 MET3
log₂ (FC)
C
Normalized fold expression of INO1
C
I
BY4743
co diploid
co S(2n-1)
co S#2
D
Normalized fold expression of KAR2
E
+ choline
- choline
WT
cho2opi3
co pox1
co pox1 S#1
co pox1 S#2
10⁰ → 10⁻⁵
absolute copy number
Chr I
Chr IV
Chr VI
Chr IX
Chr XV


Figure S6

**Figure S6. Transcript Profiling of Evolved *cho2opi3* 2n-1 Suppressors, Related to Figure 6**

(A) Scatter plot comparing the transcript profile *vs*. wild type of a *cho2opi3* mutant deprived of choline for 13 h in inositol-free medium to the slow growth signature (taken from O'Duibhir et al., 2014). Transcripts with a change in expression level with $p < 0.01$ were included.

(B) Transcript profiles of the PC-depleted parent strain and evolved suppressors *vs*. wild type, showing the 20 stress response genes (GO-BP:0006950) exhibiting the strongest increase in expression, genes involved in fatty acid and phospholipid biosynthesis, and autophagy that change more than 1.4-fold ($p < 0.01$) in at least one of the comparisons (not caused by their location on chromosome XV), and genes governing sulfur amino acid biosynthesis changing more than 1.7-fold ($p < 0.01$) in parent or suppressor *vs*. wildtype. Heatmaps show $\log_2$ fold changes vs. wild type; asterisks (*) indicate genes on chromosome XV; genes in bold contain $UAS_{INO}$.

(C and D) *INO1* (C) and *KAR2* (D) transcript levels after switching the strains indicated from SD $C^+I^+$ to the medium indicated at $OD_{600}$ 0.02 or 0.2 (*co* diploid in $C^-$), and culture for 24 h at 30 °C. Data were internally normalized to *ACT1* and expressed relative to the corresponding strain cultured in $C^+I^+$. The error bars represent SD (n=3).

(E) Growth on SD $C^{+/-}$ (30 °C for 3 d) of two *co pox1* suppressor clones obtained after 10 d incubation of *co pox1* on SD $C^-$ plates, compared to that of the strains indicated, and absolute copy numbers of chr I, IV, VI, IX and XV in *co pox1* and two independent *co pox1* suppressor clones, as determined by qPCR and FACS.

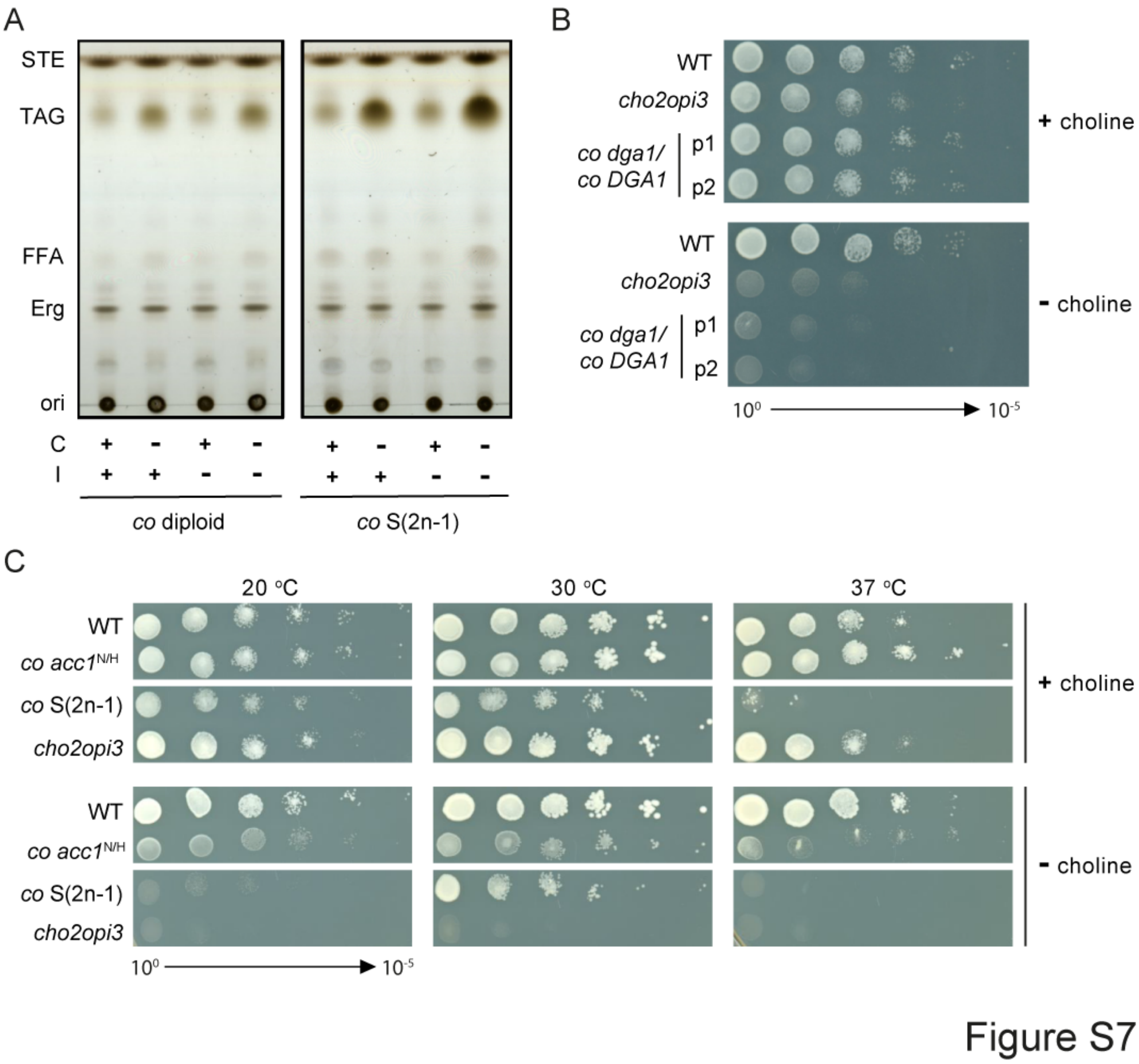


**Figure S7. FFA, *DGA1*, Related to Figure 7**

(A) TLC analysis of neutral lipids shows accumulation of free fatty acids (FFA) in *co* S(2n-1) cultured without choline; STE, sterolester.

(B) Loss of 1 copy of *DGA1* does not rescue the choline auxotrophy of the *co* diploid strain. Serial dilution experiment comparing haploid wild type and *cho2opi3* to 2 clones of heterozygous diploid *co*/*co DGA1*/*dga1* on SD $C^{+/-}$ after incubation for 3 d at 30 °C.

(C) Growth phenotype of the strains indicated after 4 d of incubation on SD $C^{+/-}$ at the temperature indicated.

**Table S1. GO-BP enrichment for transcripts changing in PC-depleted *cho2opi3* and in all 3 PC-free aneuploid *cho2opi3* suppressors tested (S#3, S#4 and S#5)**

Changes in transcript levels were used for enrichment analysis based on a fold-change cutoff of 1.7. Transcripts with decreased expression in the aneuploid *cho2opi3* suppressors encoded by chromosome XV were omitted from the analysis. GO term enrichment was considered significant with a p-value less than 0.01 (after Bonferroni correction). Enriched GO terms were summarized by the REVIGO software using a cutoff value C of 0.5 (Supek et al., 2011).

**Transcripts with increased expression: *cho2opi3 vs.* WT**

| Gene Ontology – Biological Process | Corrected p-value | Cluster frequency | Background frequency |
|---|---|---|---|
| oxidation-reduction process (GO:0055114) | 1,08E-43 | 120/420 (28.6%) | 442/6439 (6.86%) |
| generation of precursor metabolites and energy (GO:0006091) | 1,88E-33 | 71/420 (16.9%) | 193/6439 (3%) |
| ATP metabolic process (GO:0046034) | 4,03E-32 | 49/420 (11.7%) | 90/6439 (1.4%) |
| drug metabolic process (GO:0017144) | 2,53E-27 | 78/420 (18.6%) | 281/6439 (4.36%) |
| small molecule metabolic process (GO:0044281) | 2,49E-26 | 133/420 (31.7%) | 764/6439 (11.9%) |
| nucleobase-containing small molecule metabolic process (GO:0055086) | 7,39E-19 | 68/420 (16.2%) | 289/6439 (4.49%) |
| carbohydrate derivative metabolic process (GO:1901135) | 3,50E-13 | 70/420 (16.7%) | 381/6439 (5.92%) |
| carbohydrate metabolic process (GO:0005975) | 4,06E-11 | 52/420 (12.4%) | 254/6439 (3.94%) |
| transmembrane transport (GO:0055085) | 1,61E-10 | 73/420 (17.4%) | 457/6439 (7.1%) |
| response to oxidative stress (GO:0006979) | 9,49E-08 | 31/420 (7.38%) | 128/6439 (1.99%) |
| protein folding (GO:0006457) | 1,64E-06 | 28/420 (6.67%) | 119/6439 (1.85%) |
| phosphorus metabolic process (GO:0006793) | 2,11E-06 | 88/420 (21%) | 724/6439 (11.2%) |
| tricarboxylic acid metabolic process (GO:0072350) | 2,27E-06 | 14/420 (3.33%) | 31/6439 (0.481%) |
| response to abiotic stimulus (GO:0009628) | 3,06E-05 | 34/420 (8.1%) | 186/6439 (2.89%) |
| sulfur compound metabolic process (GO:0006790) | 1,11E-04 | 27/420 (6.43%) | 134/6439 (2.08%) |
| cofactor metabolic process (GO:0051186) | 1,19E-04 | 40/420 (9.52%) | 253/6439 (3.93%) |
| response to chemical (GO:0042221) | 2,68E-04 | 67/420 (16%) | 551/6439 (8.56%) |
| primary alcohol metabolic process (GO:0034308) | 3,48E-04 | 9/420 (2.14%) | 17/6439 (0.264%) |
| intron homing (GO:0006314) | 2,17E-03 | 6/420 (1.43%) | 8/6439 (0.124%) |
| cellular aldehyde metabolic process (GO:0006081) | 4,24E-03 | 14/420 (3.33%) | 52/6439 (0.808%) |

**Transcripts with decreased expression: *cho2opi3 vs.* WT**

| Gene Ontology – Biological Process | Corrected p-value | Cluster frequency | Background frequency |
|---|---|---|---|
| cellular amino acid metabolic process (GO:0006520) | 2,89E-13 | 40/233 (17.2%) | 249/6439 (3.87%) |
| ribonucleoprotein complex biogenesis (GO:0022613) | 2,40E-05 | 47/233 (20.2%) | 566/6439 (8.79%) |
| small molecule metabolic process (GO:0044281) | 2,88E-05 | 57/233 (24.5%) | 764/6439 (11.9%) |

| | | | |
|---|---|---|---|
| imidazole-containing compound metabolic process (GO:0052803) | 1,18E-04 | 6/233 (2.58%) | 9/6439 (0.14%) |
| ribosomal large subunit biogenesis (GO:0042273) | 1,10E-03 | 17/233 (7.3%) | 122/6439 (1.89%) |
| one-carbon metabolic process (GO:0006730) | 1,23E-03 | 7/233 (3%) | 18/6439 (0.28%) |
| organic acid transmembrane transport (GO:1903825) | 1,71E-03 | 12/233 (5.15%) | 64/6439 (0.994%) |
| ncRNA metabolic process (GO:0034660) | 4,98E-03 | 42/233 (18%) | 577/6439 (8.96%) |
| iron ion homeostasis (GO:0055072) | 5,73E-03 | 11/233 (4.72%) | 60/6439 (0.932%) |

**Transcripts with increased expression: aneuploid suppressors co S#3, #4, #5 *vs.* WT**

| **Gene Ontology – Biological Process** | **Corrected p-value** | **Cluster frequency** | **Background frequency** |
|---|---|---|---|
| oxidation-reduction process (GO:0055114) | 1,43E-05 | 16/49 (32.7%) | 442/6439 (6.86%) |
| sulfur amino acid biosynthetic process (GO:0000097) | 1,21E-04 | 6/49 (12.2%) | 41/6439 (0.637%) |
| small molecule metabolic process (GO:0044281) | 1,08E-03 | 18/49 (36.7%) | 764/6439 (11.9%) |
| sulfur compound metabolic process (GO:0006790) | 1,41E-03 | 8/49 (16.3%) | 134/6439 (2.08%) |
| fatty acid catabolic process (GO:0009062) | 3,19E-03 | 4/49 (8.16%) | 20/6439 (0.311%) |

**Transcripts with decreased expression: aneuploid suppressors co S#3, #4, #5 *vs.* WT**

| **Gene Ontology – Biological Process** | **Corrected p-value** | **Cluster frequency** | **Background frequency** |
|---|---|---|---|
| response to pheromone triggering conjugation with cellular fusion (GO:0000749) | 2,75E-06 | 11/106 (10.4%) | 64/6439 (0.994%) |
| cellular amino acid biosynthetic process (GO:0008652) | 2,69E-04 | 17/106 (16%) | 131/6439 (2.03%) |
| one-carbon metabolic process (GO:0006730) | 4,27E-03 | 5/106 (4.72%) | 18/6439 (0.28%) |
| small molecule metabolic process (GO:0044281) | 5,05E-03 | 29/106 (27.4%) | 764/6439 (11.9%) |

### Table S2. Primers used, Related to KEY RESOURCES TABLE and STAR Methods

| Construct | Oligos | |
|---|---|---|
| *OPI3-LEU2*[1] | Forward | 5'CAGAGCCATAAACAGCAATTGAAGACAACAAGAATAGCGTCGTAAGATGCAAGAGTTCG3' |
| | Reverse | 5'GCATAGGCTTCTAACATTATAGAATATATAGAAATAGAGCACCCCTCCTCCTTGTCAATATTA3' |
| *OPI3-LEU2* control PCR | A | 5'TGCTTCCTTGATGACCAGGGT3' |
| | D | 5'CACTTTCGCGAAGAAATTGC3' |
| *LRO1-HIS3*[1] | Forward | 5'TCTACTAAATACCGATACGAAGAAGCGTATAGTAACAGCCCGTACGCTGCAGGTCGAC3' |
| | Reverse | 5'CTTTGAAATAATACACGGATGGATAGTGAGTCAATGTCGGATCGATGAATTCGAGCTCG3' |
| *LRO1-HIS3* control PCR | A | 5'AAATTGAATGCCCAAGAAGGTG3' |
| | B | 5'CGGCAAAGTTACCCATAGCCT3' |
| | C | 5'TAATGACCATCATCGTGCTG3' |
| | D | 5'ACTGGCACAAGCCATACCTC3' |
| *DGA1-HIS3*[1] | Forward | 5'CGACAGTGGTCTATCAGGCTTGGATCTTTCACTACACTTCCGGATCCCCGGGTTAATTAA3' |
| | Reverse | 5'AGCTCCATGACTTCGATGACCACTACAGATATATTACACACGAATTCGAGCTCGTTTAAAC3' |
| *DGA1-HIS3* control PCR | A | 5'GAAGTACTTCACCACGGGGG3' |
| | D | 5'GAATCCGCGGATGGTTACAA3' |
| *POX1-HIS3*[1] | Forward | 5'TGACACATTTTAAGCCCTATATTTACGGTATTAGTTGATTCGTACGCTGCAGGTCGAC3' |
| | Reverse | 5'TCACTAGGATTTTGTAACTTTTTGTTAAAACTTGGCGACAATCGATGAATTCGAGCTCG3' |
| *POX1-HIS3* control PCR | A | 5'AATCACCGCCCATATTCTTCC3' |
| | B | 5'CGGCAAAGTTACCCATAGCCT3' |
| | C | 5'TAATGACCATCATCGTGCTG3' |
| | D | 5'CTTCATTCCAACAAGTGCCA3' |
| *CEN15*::*pGal1-CEN15-URA3*$_{kl}$ control PCR | A | 5'TCAACCAACCTCAAACTCTCAG3' |
| | B | 5'GACGTTCGTTCGACTGATGA3' |
| | C | 5'TTTGGTACGCTTCCCATCCAG3' |
| | D | 5'TGGGTTCCTTTAACGCTTC3' |
| gRNA[2] | Forward | 5'TGCGCATGTTTCGGCGTTCGAAACTTCTCCGCAGTGAAAGATAAATGATCGCCTTGATCAAT**A**$^{657039}$ACGTTTCGTTTTAGAGCTAGAAATAGCAAGTTAAAATAAGGCTAGTCCGTTATCAAC3' |
| | Reverse | 5'GTTGATAACGGACTAGCCTTATTTTAACTTGCTATTTCTAGCTCTAAAACGAAACGTT**T**ATTGATCAAGGCGATCATTTATCTTTCACTGCGGAGAAGTTTCGAACGCCGAAACATGCGCA3' |
| Repair DNA[3] | Forward | 5'TTAGAATCATCATCAAAGATCCTCAAACAGGTGCCCCAGTACCATTGCGTGCCTTGATCAAT**C**$^{657039}$ACGTTTCTGGTTATGTTATCAAAACAGAAATGTACACCGAAGTCAAGAACGCAAAA3' |
| | Reverse | 5'CTTTTGCGTTCTTGACTTCGGTGTACATTTCTGTTTTGATAACATAACCAGAAACGT**G**ATTGATCAAGGCACGCAATGGTACTGGGGCACCTGTTTGAGGATCTTTGATGATGATTCTAA3' |
| *ACC1-acc1*$^{N1446H}$ | Forward | 5'TCGTCGTGCTTATCGTGCTT3' |
| | Reverse | 5'ACAGTACGTTCAGTGAGAACAGA3' |
| | Sequencing | 5'AGGAAGATTGTCCAACTTCAAC3' |

[1] The underlined sequences correspond to nucleotides upstream (forward) and downstream (reverse complementary) of the gene to be deleted.

[2] The underlined sequences present the *ACC1* target sequence (that is followed by the PAM [NGG] on the chromosome); the flanking sequences overlap with both sides of the linearized pMEL16 backbone. The nucleotide in bold indicates the nucleotide to be edited.

[3] The nucleotide in bold indicates the edited nucleotide.
